# Specific Mitotic Events Drive Cytoskeletal Remodeling Required for Left-Right Organizer Development

**DOI:** 10.1101/2024.05.12.593765

**Authors:** Yan Wu, Yiling Lan, Favour Ononiwu, Abigail Poole, Kirsten Rasmussen, Jonah Da Silva, Abdalla Wael Shamil, Li-En Jao, Heidi Hehnly

## Abstract

Cellular proliferation is vital for tissue development, including the Left-Right Organizer (LRO), a transient organ critical for establishing the vertebrate LR body plan. This study investigates cell redistribution and the role of specific progenitor cells in LRO formation, focusing on cell lineage and behavior. Using zebrafish as a model, we mapped all mitotic events in Kupffer’s Vesicle (KV), revealing an FGF-dependent, anteriorly enriched mitotic pattern. With a KV-specific fluorescent microtubule (MT) line, we observed that mitotic spindles align along the KV’s longest axis until the rosette stage, spindles that form after spin, and are excluded from KV. Early aligned spindles assemble cytokinetic bridges that point MT bundles toward a tight junction where a rosette will initially form. Post-abscission, repurposed MT bundles remain targeted at the rosette center, facilitating actin recruitment. Additional cells, both cytokinetic and non-cytokinetic, are incorporated into the rosette, repurposing or assembling MT bundles before actin recruitment. These findings show that initial divisions are crucial for rosette assembly, MT patterning, and actin remodeling during KV development.

## Introduction

The Left-Right Organizer (LRO) is one of the earliest ciliated organs formed during vertebrate development. It is a small and yet important structure—e.g., the node in mice and Kupffer’s vesicle (KV) in zebrafish—that only exists transiently during early embryogenesis to help establish the left-right patterning of the body plan. The LRO initiates the development of left-right asymmetry through its cilia-driven, directional fluid flow that induces asymmetric gene expression (summarized in ^1–4^). Compromised formation of LRO or its ciliary motility (e.g., mutations in the axonemal dynein motor ^5–7^) would lead to LR patterning defects.

Previous studies using the mouse and zebrafish models (**Fig 1A**, reviewed for fish in ^1,3^, study in mice ^8^) have shown that LRO is formed through the organization of the LRO progenitor cells (e.g., dorsal forerunner cells in zebrafish) into rosette-like structures before displaying epithelial-like characteristics and initiating lumen formation. Using in vitro three-dimensional (3D) cultured Madin-Darby Canine Kidney (MDCK) cells suspended in extracellular matrix ^9–12^, lumen formation has been further shown to be preceded by cell division, followed by the establishment of apical polarity and cell-cell contacts adjacent to the cytokinetic bridge. However, those in vitro 3D cell culture models do not recapitulate the mesenchymal to epithelial transition, nor the dynamic changes in cell shape and expression profiles during LRO formation in a living organism. In addition, without the in vivo biological context, different in vitro 3D models often lead to different conclusions. For example, the MDCK ^13^ and Inner Medullary Collecting Duct (IMCD)^14^ models showed a different timing of cilia formation. Therefore, despite the insights gained from different in vitro 3D cell culture models, the cellular mechanisms and signals underlying the formation and maintenance of the LRO remain unresolved.

**Figure 1.**
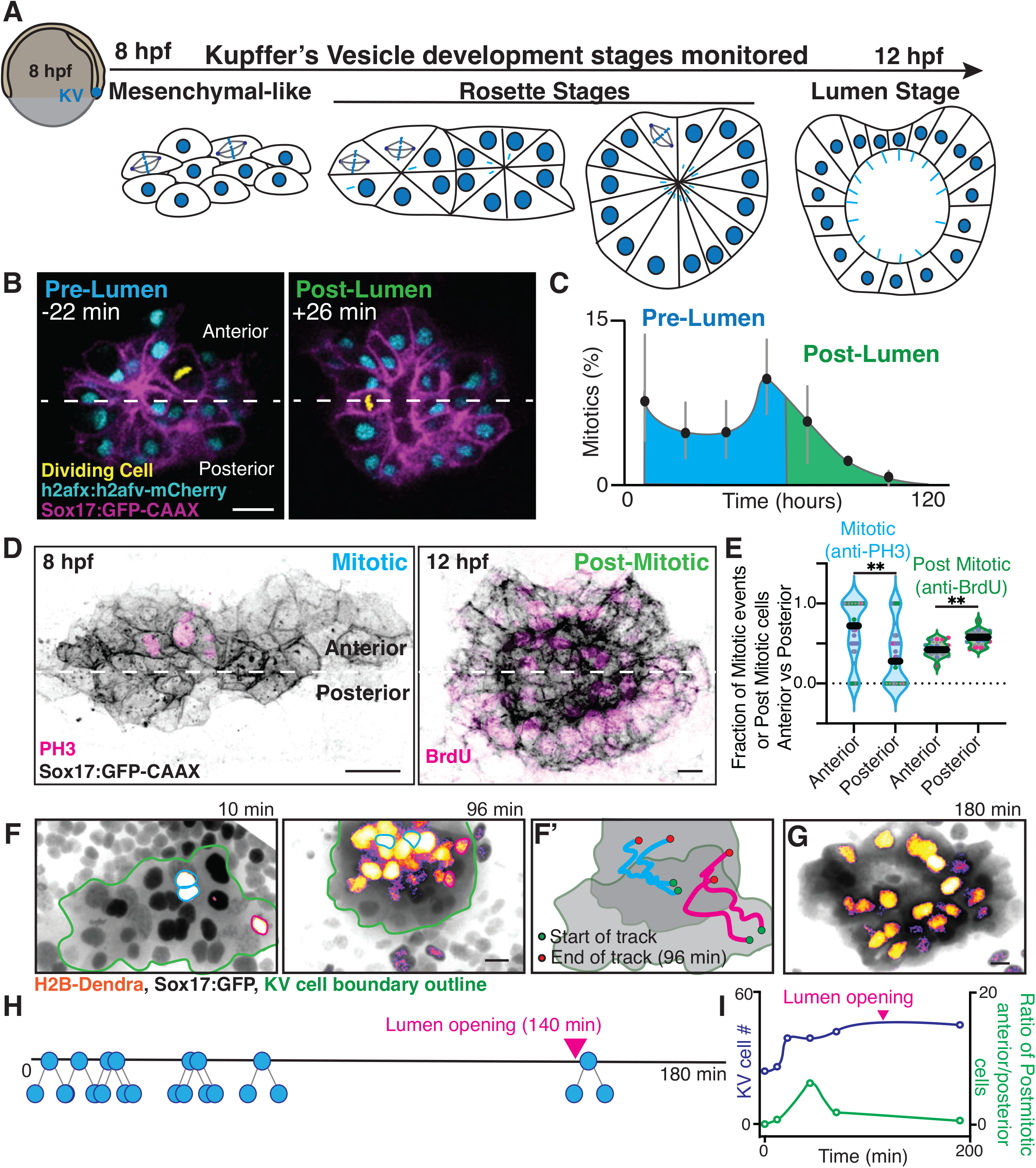
Identification of an anteriorly positioned and pre-lumen enriched Kupffer’s Vesicle (KV) mitotic events and their post-mitotic distribution map. **(A)** Model depicting KV cells transitioning from a small population, likely proliferative, then organize into rosette structures then transition to a cyst of ciliated (light blue) cells surrounding a fluid filled lumen. **(B)** Representative confocal volumetric projections from live KV development video (refer to **Video S1**). Pre-KV lumen time point (-22 min before lumen opening) and post-lumen (26 min past lumen opening) shown. KV cell membranes marked by Sox17:GFP-CAAX (magenta) and nuclei with h2afx:h2afv-mCherry (non-mitotic, cyan; mitotic, yellow). Scale bar, 10 μm. **(C)** Percentage of KV cells that are mitotic across a 120 min video time course (n=3 embryos ±SEM). **(D)** Confocal projections of mitotic (Phospho-Histone H3, PH3, magenta) and post-mitotic (BrdU, magenta) events at pre lumen (8 hpf) and post lumen (12 hpf). KV cell membranes marked with Sox17:GFP-CAAX (inverted gray). Scale bar, 10 μm. **(E)** Fraction of PH3 and BrdU-positive cells in the anterior vs posterior regions of the KV (n=4 clutches, 36 embryos; and n=3 clutches, n=17 embryos respectively). Each point represents an embryo, the point colors represent clutch, n=3 clutches, **p<0.01. **(F-G)** Confocal projections of H2B-Dendra photoconversion (non-photoconverted black nuclei, photo-converted in fire-LUT). Cyan and magenta outlines mark mitotic photoconverted event at 10 min. A total of 8 mitotic events were photoconverted. Daughter cell positioning at (**F**) 96 min and (**G**) 180 min shown. **(F’)** Trajectory of marked mitotic events from division completion to 96 min. KV cell boundary, green outline. Scale bar, 10 μm. **(H)** Temporal schematic of KV mitotic event for representative embryo in (**F**), daughter cells did not divide again. **(I)** KV cell number (blue) and ratio of post mitotic anterior positioned or posterior positioned cells were calculated over time from acquisition in (**F-G**). Statistical results detailed in **Table S1**.

To understand how a functional LRO is formed in a living organism—from a small group of precursor cells to a fluid-filled sac lined with both motile and non-motile ciliated cells ^15,16^, we studied the development of the KV at single-cell resolution using a series of novel transgenic zebrafish lines and cell biological techniques, including high-resolution live microscopy, lineage tracing, laser ablation, and pharmacological treatments. Our study shows that FGF signaling-mediated early divisions of KV precursor cells are pivotal for proper KV development as the progeny of these early division events constitute a proportion of cells in the mature KV. Our data also suggest that apical microtubule (MT) bundles—originated from the daughter cells of the early KV precursors—facilitate actin recruitment that is crucial for KV development.

## Results

### Identification of an anteriorly positioned and pre-lumen enriched KV mitotic events and their post-mitotic distribution map

To follow how the KV is formed at single-cell resolution in a living vertebrate, a Tg(Sox17:GFP-CAAX; h2afx:h2afv-mCherry) was used, where the KV progenitor cells, i.e., the dorsal forerunner cells (DFCs), were labeled by membrane-bound GFP while all histones were labeled by the red fluorescent protein mCherry. We followed the movement and division of DFCs by time-lapse microscopy starting at approximately 8 hours post fertilization (hpf) when 15–30 DFCs have formed (**Fig 1A, 1B, Video S1**). The KV transitions from approximately 15 DFCs that then expand in number and organize into rosette configurations before transitioning to a lumen (**Fig 1A**). Mitotic events predominantly occurred in the anterior region of the future KV before lumen opening (**Fig 1B-C, Fig S1A-B, Video S1**). The same conclusion was also reached when fixed Sox17:GFP-CAAX embryos were analyzed by mitotic cell marker, anti-phospho-histone 3 (PH3) staining at 8 hpf, where we detected a significant fraction of mitotic events in the anterior zone (**Fig 1D, 1E**) with an enrichment to the left anterior quadrant (**Fig S1C**).

Next, we investigated the distribution of post-mitotic daughter cells from KV precursors in the fully formed KV. For this purpose, embryos were incubated with BrdU from 70% epiboly to 12 hpf to label cells that had undergone cell division before KV completion at 12 hpf. We observed that these BrdU-positive (post-mitotic) cells were distributed throughout the KV, with a significantly higher concentration in the posterior region (**Fig 1D-E, Fig S1D**).

To determine specifically how KV progenitor cells and their progeny reposition following cell division, we performed cell lineage tracing analysis of H2B-Dendra expressing, Sox17:GFP marked KV precursor cells (**Fig 1F-G**). In this experiment, individual H2B-Dendra-positive cells (green) were photoconverted at metaphase and the trajectories of the resulting daughter cells (red) were followed throughout the KV developmental stages until lumen expansion (**Fig 1A**, **Fig 1F-G**). We found that the daughter cells of a given KV precursor stay together and move within the KV boundary after division (representative events shown in **Fig 1F’** and **Fig S1E**). Among 8 photoconverted metaphase cells and their progeny were followed, all the daughter cells did not divide again during KV development, indicating that at least 50% of KV progenitor cells divide once and only once after 8 hpf (**Fig 1H**). We found that most of the post-mitotic cells were organized in the anterior region of the KV until the rosette stage at the 120 min time point, and then redistributed to the posterior region of the KV during lumen formation (180 min, **Fig 1H, 1I**). When we determined the ratio of anteriorly versus posteriorly positioned post-mitotic cells, this ratio started to approach 1 right before lumen opening and after the cell number in the KV had reached its maximum (**Fig 1I**). Together these results indicate that KV progenitor cells divide preferentially in the anterior region of the future KV, and the resulting post-mitotic cells were incorporated throughout the mature KV with more cells in the posterior than in the anterior regions.

### An anterior/posterior FGF signaling gradient is required for anterior zones of mitotic activity

Since the early mitotic events of KV precursors took place predominantly anterior to the location of the future KV (**Fig 1**), we hypothesized that an anteroposterior signal(s) coordinates the cell division and the later LRO development. To test this, we individually inhibited the major signaling pathways that had been implicated in LRO development ^17–19^ using well-established pharmacological inhibitors, and then determined whether KV development is affected (**Fig 2A, Fig S2A-E**). We found that blocking fibroblast growth factor (FGF) signaling, but not Wnt or Sonic hedgehog (Shh) signaling, attenuated the division of KV precursors through PH3 staining at 8 hpf (**Fig 2B, 2D**) and caused KV cells to not expand in number at pre-lumenal to lumenal stages (**Fig S2E**). Inhibiting FGF signaling also disrupted spatially concentrated mitotic events in the anterior quadrant of the developing KV and caused severe KV developmental defects (**Fig 2B, 2E-F, S2A-D**). Specifically, only FGF inhibition demonstrated a significant defect in KV development measured by lumen area (**Fig 2B, 2F, S2A-C**). These findings suggest that an anteroposterior FGF signaling gradient is required for the mitotic activity in the anterior region of the developing KV.

**Figure 2.**
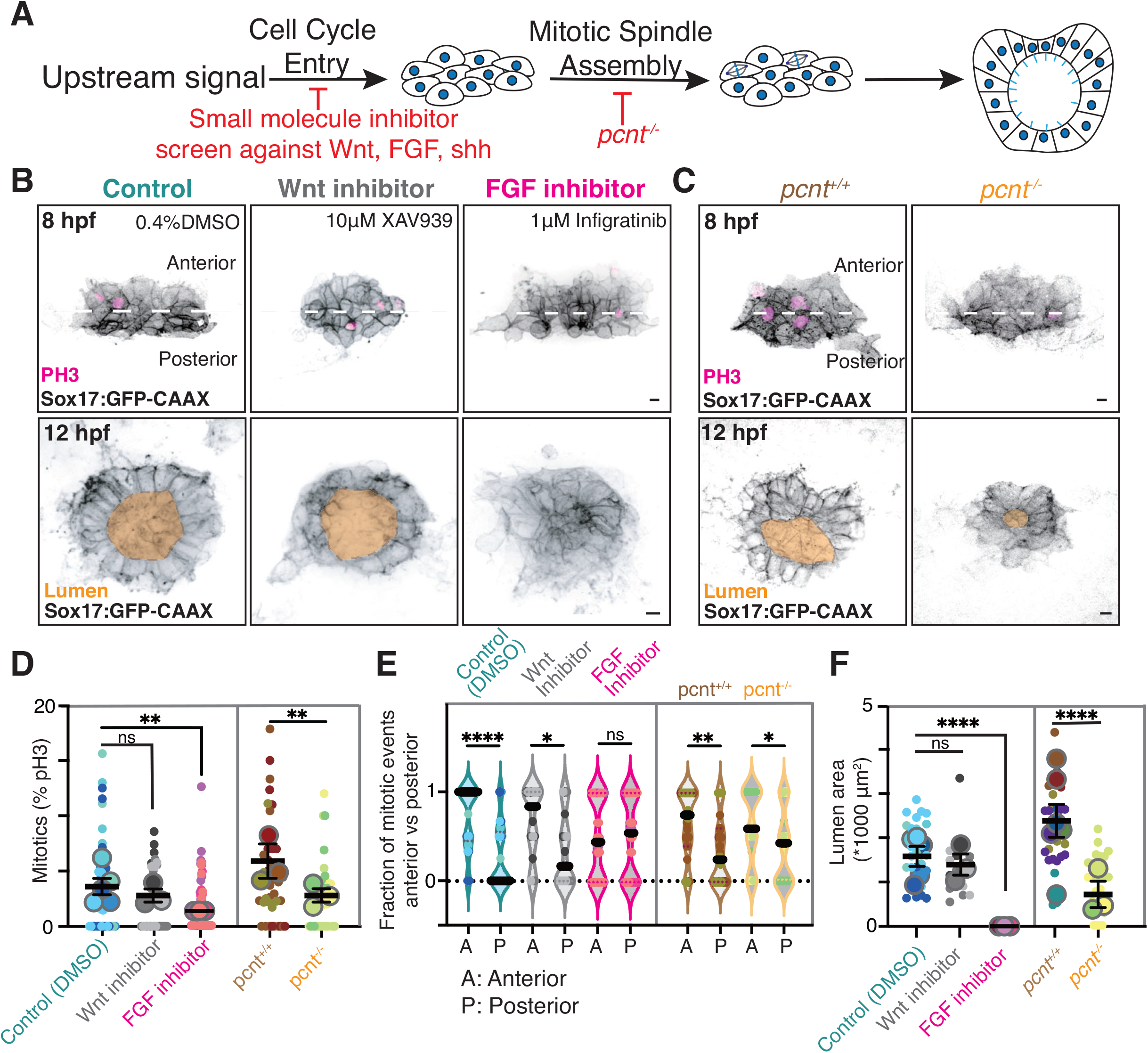
An anterior/posterior FGF signaling gradient is required for anterior zones of mitotic activity. **(A)** Model illustrating experimental approach to identify whether a signal influences mitotic events and how those events contribute to KV development. **(B-C)** Confocal projections of pre-lumen (8hpf, top) and post-lumen (12 hpf, bottom) KVs treated with DMSO, 10 μM XAV939 (Wnt inhibitor), and 1μM infigratinib (FGF inhibitor) (**B**) or lacking Pericentrin (*pcnt*^-/-^) compared with wild-type control (*pcnt*^+/+^) (**C**). KV cell membranes marked with Sox17:GFP-CAAX (inverted gray), mitotic events with PH3 (magenta). Lumens highlighted in gold. Scale bar, 10 μM. **(D-F)** Percent of mitotic KV cells (**D**), anterior vs posterior positioned mitotic events (**E**), and lumen area (**F**) were measured for n>33 embryos (small dots represent embryos, **D-F**) across n≥3 clutches (large dots represent clutch, **D, F**) per condition. Refer to Figure S2. Graphs are scatter plots with mean of clutches shown ±SEM (**D, F**) or a violin plot (**E**). DMSO, 10 μM XAV939, 1 μM infigratinib, *pcnt*^+/+^ and *pcnt*^-/-^ shown. *p<0.05, **p<0.01, ***p<0.005, ****p<0.0001, ns, not significant. Statistical results detailed in **Table S1**.

To determine whether loss of the anteriorly biased mitotic events is sufficient to disrupt proper KV formation at the later stage, we disrupted mitotic events independent of FGF signaling using pericentrin maternal-zygotic mutant embryos (*pcnt*^-/-^, ^20^) and assessed KV development. Pericentrin is a centrosomal protein known to regulate spindle assembly, and loss of pericentrin can result in a decrease in mitotic entry and failure to complete mitosis ^21–25^. We found that the overall cell division (**Fig 2C-D**) and the anterior enrichment of mitotic events (**Fig 2E**) of KV precursors decreased significantly in maternal-zygotic *pcnt*^-/-^ embryos. This decrease in anteriorly positioned mitotic events was also associated with a disruption in KV development, as demonstrated by a significant reduction in the lumen area at 12 hpf (**Fig 2C, 2F**). Together, these findings suggest that potential anteroposterior FGF signaling regulates the early, anteriorly biased KV precursor divisions, which are required for subsequent KV formation.

### Early KV mitotic events hold greater significance to KV development compared to later KV mitotic events

To determine the role and relevance of individual KV mitotic events, we employed laser ablation to selectively remove mitotic events (**Fig 3A-B**). Temporal characterization of mitotic events involved six ablation conditions, encompassing two control scenarios—no ablation and ablation of non-mitotic cells (3 to 8 cells). The first experimental ablation comparison was with ablating all mitotic events compared to no ablation controls (**Fig 3C, 3E, 3G**). Then specific temporal ablation patterns were performed and compared (**Fig 3D, 3F, 3H**): (1) ablating the first 4 mitotic events when fewer than 20 KV precursor cells were present (Condition 1); (2) ablating 4 mitotic events when more than 20 KV precursor cells were present (Condition 2); and (3) ablating all mitotic events when more than 20 KV precursor cells were present (Condition 3). During video acquisition, mitotic cells were ablated by positioning a region of interest (ROI) over the metaphase plate, distinguished by h2afx:h2afv-mCherry, and delivering a 355 nm pulsed laser within the ROI (**Fig 3B**). Monitoring encompassed all cells, including the cell with the ablated plate, and revealed no evident apoptotic events in the non-ablated cells. The ablated cells failed to complete mitosis and were ultimately extruded from the developing KV.

**Figure 3.**
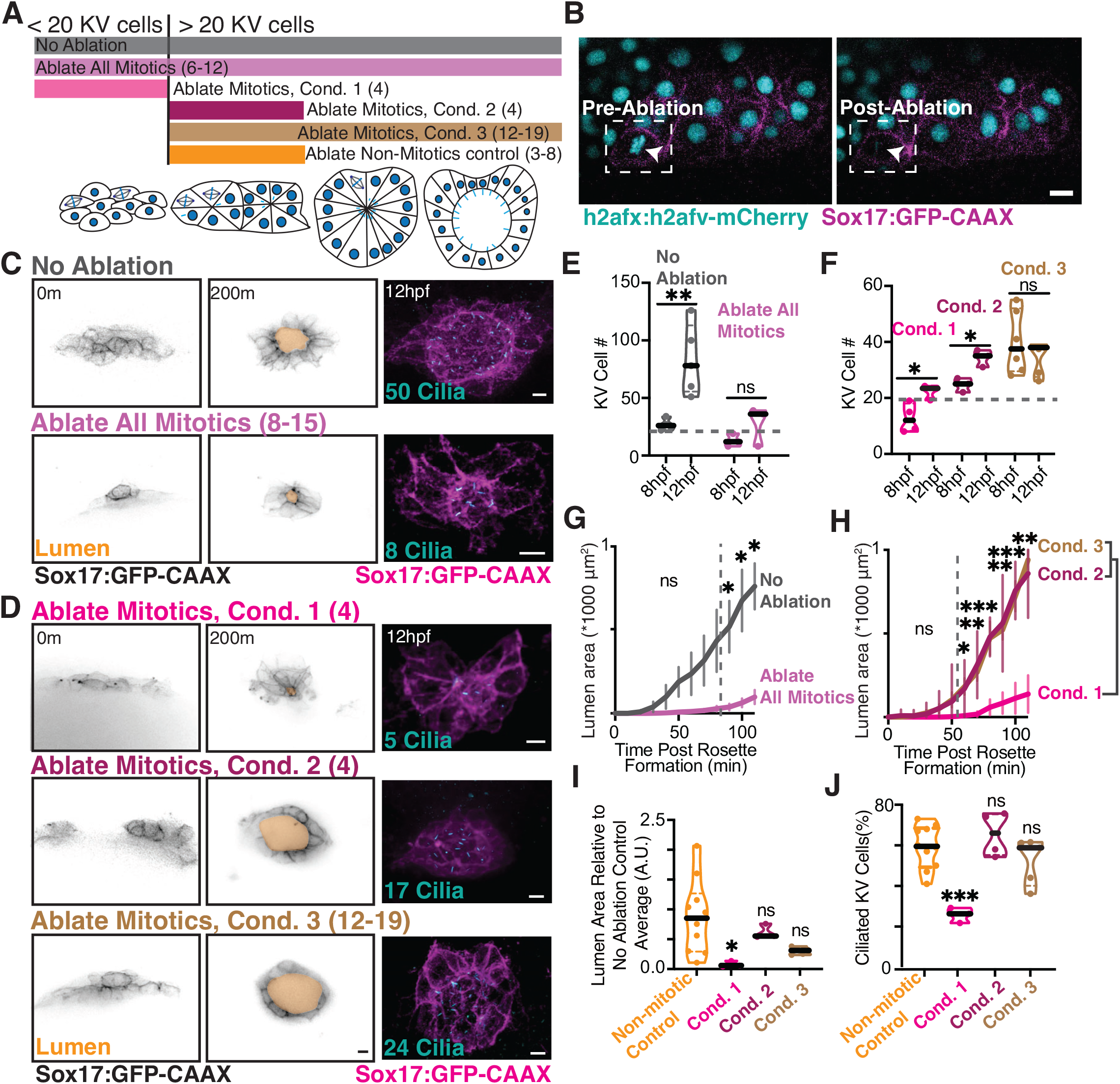
Early KV mitotic events hold greater significance to KV development compared to later KV mitotic events. **(A)** Model illustrating laser ablation conditions during KV development. **(B)** Representative images of KV ablation experiment showing a pre-ablation and post-ablation cell. White arrow highlights KV cell ablated. KV cell membranes marked by Sox17:GFP-CAAX (magenta) and nuclei with h2afx:h2afv-mCherry (cyan). Scale bar, 10 μm. **(C, D)** Confocal projections taken from a video acquisition (0-200m, left, Sox17:GFP-CAAX, inverted gray; lumen, gold, Refer to **Video S2**). The embryo was then fixed at 6SS (12 hpf) and stained for cilia (acetylated tubulin, cyan, right; Sox17:GFP-CAAX in magenta). Number of cilia in each representative image labeled. Scale bar, 10 μm. **(E, F)** Truncated violin plot depicting the number of KV cells at 8 hpf and 12 hpf for no ablation and ablating all mitotics (**E**) or ablation conditions 1, 2, or 3 (**F**) across n≥3 embryos (each point represents a single embryo). Median denoted with black line and early/late temporal zone threshold (20 cells) denoted with grey dashed line. (**G, H**) Lumen area was measured over time for over 100 min for no ablation (control) and ablating all mitotics (**G**) or ablation conditions 1, 2 or 3 (**H**). n≥3 ±SEM embryos measured per condition. Unpaired two-tailed t-tests in (**G**), One way ANOVA with Dunnett’s multiple comparison in (**H**). *p<0.05, **p<0.01, ***p<0.005, ns, not significant. (**I, J**) Truncated violin plots depicting KV lumen area (12 hpf, **I**) and percentage of ciliated KV cells (**J**). n≥3 embryos at each condition. One way ANOVA with Dunnett’s multiple comparison to non-mitotic ablation controls. *p<0.05, ***p<0.005, ns, not significant. Statistical results detailed in **Table S1**.

Our studies indicate that the first 4 KV divisions are crucial for KV development, while later divisions near lumen formation are not essential (**Fig 3C-I**). KVs did not significantly increase in cell number between 8 hpf and 12 hpf when all mitotic events were ablated or when all mitotic events were ablated with more than 20 KV precursor cells present (**Fig 3E-F**), suggesting the importance of mitotic events in KV cell number expansion during development. Significant defects in lumen formation kinetics (**Fig 3C-D, 3G-H, Video S2**) and the final lumen area (**Fig 3I**) occurred when all mitotic events or only the first 4 events were ablated (Condition 1), but not when later mitotic events were ablated (Condition 2 and 3), compared to non-mitotic cell ablation control (**Fig 3I**). Embryos subjected to ablation and monitoring for lumen formation kinetics (**Fig 3C-D**) were fixed at 12 hpf and immunostained for cilia. Like lumen formation defects, significant reductions in the percentage of ciliated cells were observed only when ablating all mitotic events or the first 4 events (**Fig 3C-D, 3J**). These findings underscore the pivotal role of early KV mitotic events in orchestrating both KV development and overall KV ciliogenesis.

### KV spindles stably align along KVs longest axis until the KV starts rounding, then spindles spin and are extruded

Proper spindle orientation during mitosis is important for tissue morphogenesis ^26,27^. Given the significance of early mitotic events in KV development (**Fig 3**), we aimed to understand the relationship between spindle positioning and division timing by analyzing the spindle behaviors in the KV precursor cells over time. To specifically label MTs in KV precursors, we adapted a similar strategy we have used before ^28^ to generate a Tg(Sox17:EMTB-3xGFP) transgenic line, where the MTs in KV precursors were labeled by GFP tagging the MT-binding domain from the protein ensconsin. This transgenic line allowed us to discern changes in intracellular MT patterning during KV development in live embryos for the first time (**Fig 4A**). Through a series of time-lapse imaging analysis, we found that the spindles of the dividing KV precursors formed and positioned primarily along the long axis of the developing KV in the pre-lumenal stage (**Fig 4A**, 8 min, **Video S3**). As the KV precursors were incorporated to form the rosette structure, the MT bundles started to concentrate at cytokinetic bridges near the rosette center (**Fig 4A**, 97 min, **Video S3**). Strikingly, as the KV structure went from a structure that had a long and short axis (**Fig 4A**, 8 min) to a structure that was more rounded (**Fig 4A**, 97 to 163 min), the spindles that formed started to rotate or “spin” and were subsequently extruded from the KV (**Fig 4A**, 163 min, **Fig 4B**, **Video S3**). These findings indicate that the rosette was formed by the daughter cells of the early-dividing KV precursors. In contrast, daughter cells from the later cell divisions were excluded from the KV. These results also explain why the later, but not the early, division events are dispensable for KV formation (**Fig 3**). To our knowledge, this is also the first evidence to show that cell extrusion is part of the developmental program of KV formation, akin to the cell extrusion process that has been documented in other developmental contexts such as the generation and maintenance of the tight epithelial barrier ^29–32^.

**Figure 4.**
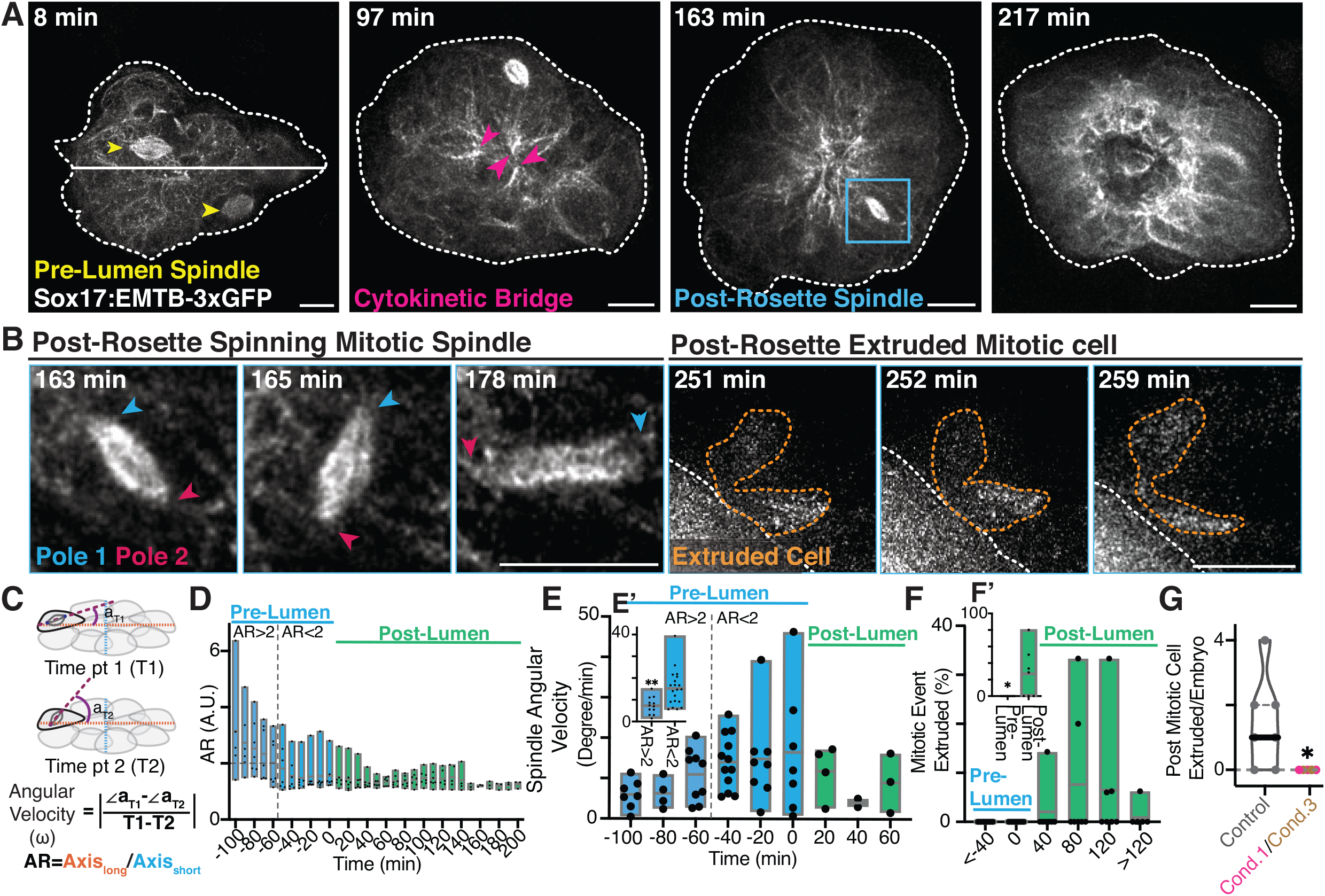
KV spindles stably align along KVs longest axis until the KV starts rounding, then spindles spin and are extruded. (**A, B**) Time-lapse video of MT organization during KV formation (Sox17: EMTB-3xGFP, gray). KV are outlined (white dashed line). Highlighted are spindles (yellow arrows) and cytokinetic bridges (pink arrows). The blue boxed region is magnified in (**B**) showcasing a spinning mitotic spindle that becomes extruded. Mitotic spindle poles noted with blue and pink arrowheads. Extruded cytokinetic KV cell post spinning event outlined (orange dashed line). Scale bars, 10 μm. Refer to **Video S3**. (**C**) Methods to measure KV aspect ratio (A.R.) and the angular velocity of the spindle (ω∠a), illustrated in model. (**D, E**) Floating bar graph depicting the maximum, minimum and mean of KV A.R. (**D**) and mean angular velocity (**E**) over 20 min binned time periods relative to lumen formation. The gray dashed line marks when A.R. equals 2. n=7 embryos measured across 6 clutches. (**E’**) Angular velocity were compared when A.R. was <2 or >2 from data sets in (**E**). 32 spindle poles from n=7 embryos across 6 clutches were monitored. Unpaired two-tailed t-tests, **p<0.01. **(F)** Floating bar graph depicting the maximum, minimum and mean of the percentage of mitotic events with post mitotic cell extruded at various time bin (each 40 min) relative to lumen formation. n=7 embryos from 6 clutches. (**F**’) Percentage of mitotic events with post mitotic cell extruded pre-versus post-lumen formation. n=7 embryos from 6 clutches. Unpaired two-tailed t-tests, *p<0.05, (G) Truncated violin plot depicting the frequency of mitotic events with post mitotic cell extruded in no ablation controls or mitotic ablated condition 1 and 3. n>6 embryos measured over 6 clutches. Unpaired two-tailed t-tests, *p<0.05. Refer to **Video S4**. Statistical results detailed in **Table S1**.

To characterize the robustness of these observed events, the aspect ratio (A. R.), defined as the longest axis over the shortest axis of the KV (**Fig 4C**), was measured during KV development (**Fig 4D**) and compared to the degree of spindle angle changes per minute (**Fig 4C, 4E**). Time was normalized to lumen opening so that negative time values are pre-lumen formation and positive values are post-lumen formation (**Fig 4D-4F**). We identified that as the aspect ratio of the KV approached a mean value of less than 2 (dashed line, **Fig 4D**), that spindles started to spin which relates to an increased rate of rotation (angular velocity, **Fig 4E**, direct comparison between A.R. less than 2 and greater than 2 in **4E’**). A population of these mitotic events then become extruded once the lumen starts to open (**Fig 4F**, direct comparison between A.R. less than 2 and greater than 2 in **4F’**). When we ablated the first four mitotic events (condition 1) or ablated all mitotic events post-reaching 20 KV progenitor cells (condition 3, refer to **Fig 3A, 4G, Video S3 and S4**), we observed a loss of post-mitotic extrusion events. Interestingly, even when we only ablated the first 4 mitotic events, no later extrusion events occurred, suggesting that extrusion may be a response to cell packing. This suggests that if early events are alleviated, later division events can still be incorporated into the KV, but the KV doesn’t develop appropriately (**Fig 3**).

### KV progenitor cells position cytokinetic bridges towards tjp1 marked tight junctions where they maintain cytokinetic bridge derived MT bundles post bridge cleavage

The findings in **Fig 4A** demonstrate that cytokinetic bridges are organized toward the center of a KV cell rosette. However, the coordination of this process with other temporally regulated cellular events, such as tight junction organization and apical actin recruitment, remains unclear. To investigate this, we utilized TgKI (tjp1atdTomato); Sox17:EMTB-3xGFP embryos to label the tight junction marker, tjp1, at its endogenous locus ^33^ while simultaneously marking MTs in KV cells. In these embryos, tjp1-labeled tight junctions were assembled prior to division events. Subsequently, early KV divisions positioned their cytokinetic bridges toward a tjp1-labeled junction (**Fig 5A, 5B**), where MT bundles derived from the cytokinetic bridge were maintained post-bridge cleavage (**Fig 5C**, arrow at 2m indicating midbody loss). This finding contrasts with studies in human cells grown in culture, which do not assemble into 3D structures, where MT enrichment at bridges diminishes as abscission completes ^34–37^. This suggests that the assembly of a 3D tissue may impose additional requirements on these MT-based structures as modeled in **Fig 5D**.

**Figure 5:**
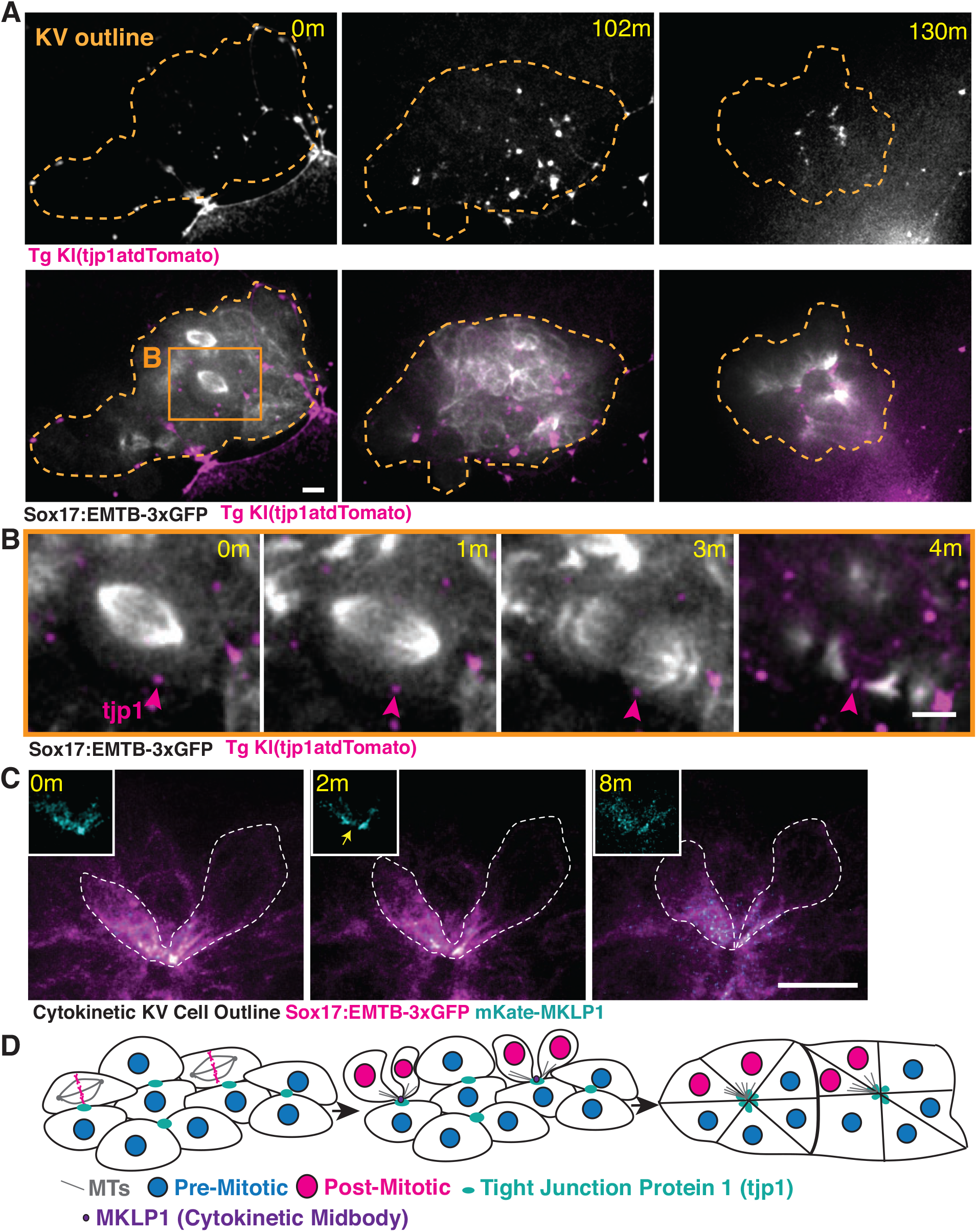
KV progenitor cells position cytokinetic bridges towards tjp1 marked tight junctions where they maintain cytokinetic bridge derived MT bundles post bridge cleavage. **(A-B)** Time-lapse image of MT (Sox17:EMTB-3xGFP, gray) and tjp1 (TgKI (tjp1atdTomato, magenta)) distribution in KV mitotic cells. Time-lapse image of metaphase cell in (**A**) highlighted by orange box is magnified in (**B**). Scale bars, 5 μm. **(C)** Time-lapse image of an abscission event (yellow arrow) with a cytokinetic bridge and midbody labeled with mKate-MKLP1 (cyan) and MT (EMTB-3xGFP, magenta). Insets show MKLP1 (cyan) decorating a midbody (0 min), bridge abscission (yellow arrow, 2 min), and midbody loss at 8 min. Scale bar, 10 μm. **(D)** Working model depicting mitotic events affecting midbody distribution and tight junction formation during KV development.

### Converging cytokinetic cells initiate actin recruitment at forming rosette centers

To examine the temporal and spatial dynamics of actin in relation to MTs, we injected Sox17:EMTB-3xGFP embryos with Lifeact-mRuby mRNA (**Fig 6A**). When imaging KV conditions with fewer than 20 cells, the first cytokinetic bridge was found to accumulate actin (magenta arrow, **Fig 6B**, **Video S5**) and serves as the initial site for rosette formation. A second cytokinetic bridge, also recruiting actin, can move towards the rosette, allowing the daughter cells connected by this bridge to integrate into the developing structure (blue arrow, **Fig. 6B**). Surrounding cells, which are not connected by cytokinetic bridges, can then assemble into the rosette, organizing MT bundles toward its center (non-division-derived MT bundle, NDMT, **Fig 6B**, modeled in **Fig 6C**).

**Figure 6:**
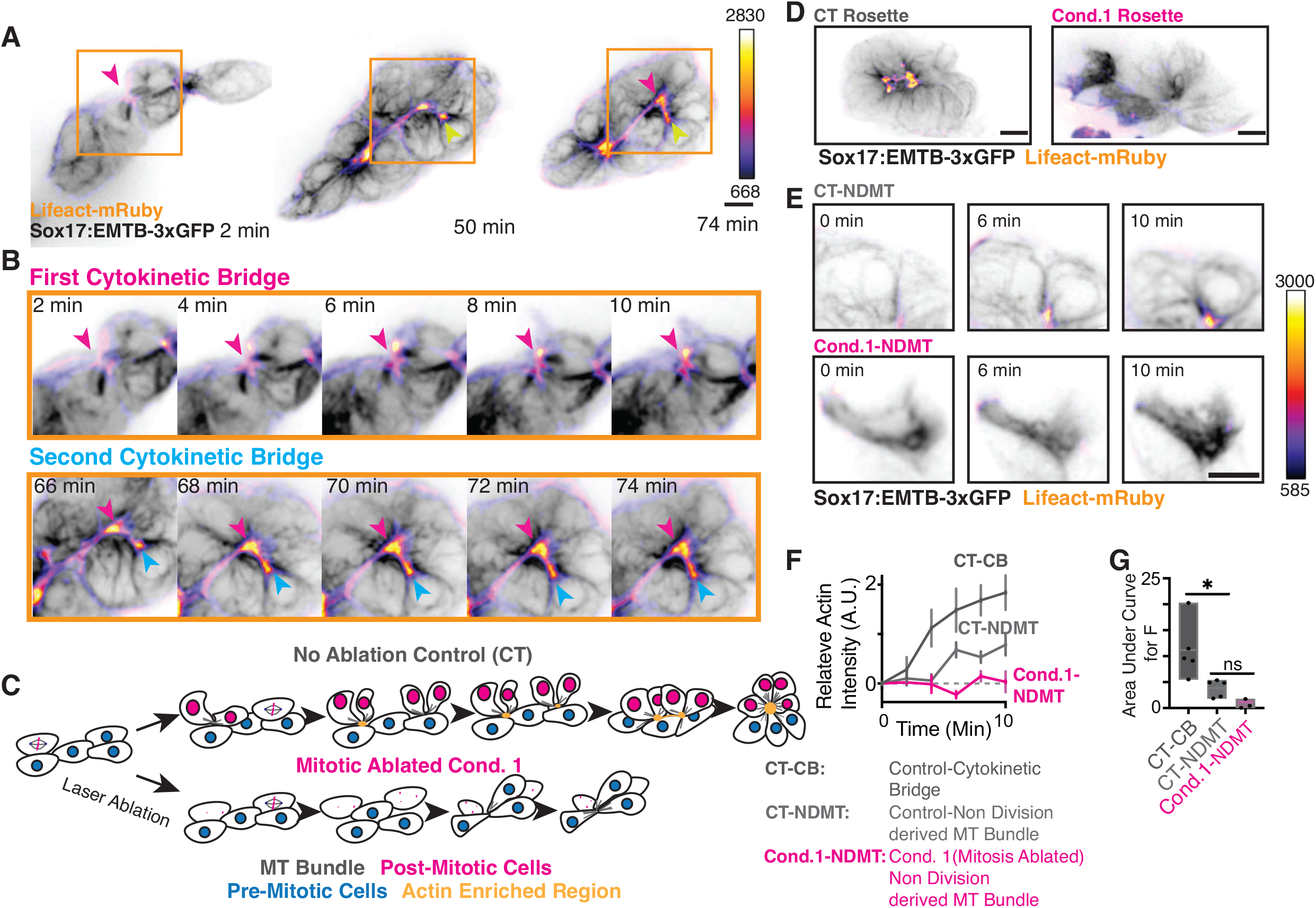
Converging cytokinetic cells initiate actin recruitment at forming rosette centers. **(A-B)** Time-lapse video of actin (Lifeact-mRuby, FireLUT) and MTs (Sox17:EMTB-3xGFP, inverted Gray). The region denoted by orange box in **(A)** is magnified 1.5X in **(B)** first cytokinetic bridge is marked (magenta arrow) in relation to second cytokinetic bridge (blue arrow). Scale bar, 10 μm, calibration bar, Lifeact-mRuby fluorescent intensity values. Refer to **Video S5.** **(C)** Testable model for how KV forms MT bundles and actin enriches in no ablation control compared to mitotic ablation conditions. **(D)** Representative confocal volumetric projections of KVs depicting MTs (Sox17:EMTB-3xGFP, inverted gray) and actin (Lifeact-mRuby, FireLUT) of no ablation control (CT, Left) or mitotic ablated condition 1 (Cond.1, right) at rosette stage. Scale bar, 10 μm. **(E)** Time lapse images of non-division-derived MT bundle formation (Sox17:EMTB-3xGFP, inverted gray) and actin (Lifeact-mRuby, FireLUT) recruitment at tips of the bundles from no ablation control condition (CT-NDMT, Top) and mitotic ablated condition 1 (Cond.1-NDMT, bottom). Scale bar, 10 μm. Calibration bar on the right denotes Lifeact-mRuby intensity. **(F-G)** Line chart depicting the relative actin intensity 10 min (**F**) or floating bar graph depicting area under curve (**G**) for relative actin intensity over 10 min interval featured in (**F**). One way ANOVA with Dunnett’s multiple comparison, *p<0.05, ns, not significant, n≥3 for each condition over ≥2 clutches. Statistical results detailed in **Table S1**.

To determine whether the initial mitotic events are required for actin recruitment to rosette centers, we ablated the first four divisions (condition 1, **Fig 3A**) and compared them to control, non-ablated conditions (**Fig 6C**). Under ablated conditions, actin was unable to accumulate at rosette centers that had accumulated non-division-derived MT bundles (NDMT, **Fig 6D**, right) compared to controls (**Fig 6D**, left). Actin recruitment to MT bundles across three conditions: 1) cytokinetic bridge-derived MT bundles (CT-CB), 2) non-division-derived MT bundles (CT-NDMT), and 3) NDMT from condition 1 ablated cells (Cond.1-NDMT). While non-division-derived MT bundles (CT-NDMT) can recruit actin (**Fig 6F**), the recruitment is significantly less compared to cytokinetic bridge-derived MT bundles over a 10-minute period (CT-CB, **Fig 6F, 6G**). However, ablating the first four mitotic cells does not significantly decrease the ability of non-division-derived MT bundles (Cond.1-NDMT) to recruit actin (**Fig 6E-G**). This suggests that while the properties of cytokinetic bridge-derived MT bundles are associated with increased actin recruitment compared to non-division-associated MT bundles, post-mitotic cells do not influence the ability of neighboring rosette cells with non-division-derived MT bundles to assemble actin.

### Cytokinetic bridge derived MT bundles are required for actin recruitment at rosette during lumen formation

To further investigate whether cytokinetic bridge-derived MT bundles are necessary for actin recruitment during the rosette stage, we conducted targeted ablations on cytokinetic bridges and non-division-derived MT (NDMT) bundles prior to the expected time of actin recruitment, then measured actin accumulation and MT recovery (**Fig 7A**). For the cytokinetic bridge ablations, we ablated one side of the bridge adjacent to where the midbody should form (CB-A), while leaving the other side unablated (CB-NA) (**Fig 7A, 7B**). We found that actin recruitment on the ablated side was significantly diminished over a 10-minute period, while the non-ablated side retained its ability to recruit actin (**Fig 7B, 7E, S3B**). When comparing the ability of MTs to recover after cytokinetic bridge ablation (CB-A) to ablated non-division-derived MT bundles (AB-NDMT), we found that MTs were unable to recover under CB-A conditions, whereas they did recover under AB-NDMT conditions (**Fig 7B-7D, 7F, S3C**). This suggests that the properties of MT bundles in the rosette differ between those derived from cytokinetic bridges and those originating from non-mitotic cells. When examining whether ablated NDMT bundles could recruit actin after 10 minutes, we found they were unable to do so compared to control non-ablated NDMT conditions (CT-NDMT, **Fig 7G, S3D**). These findings suggest that while the ability of MTs to recover varies between cytokinetic bridge-derived MT bundles and non-division-derived MT bundles (**Fig 7F**), both MT bundled structures are necessary for actin accumulation (**Fig 7E, 7G**), with cytokinetic bridge-derived bundles recruiting actin to a greater extent (**Fig 6F, G**).

**Figure 7.**
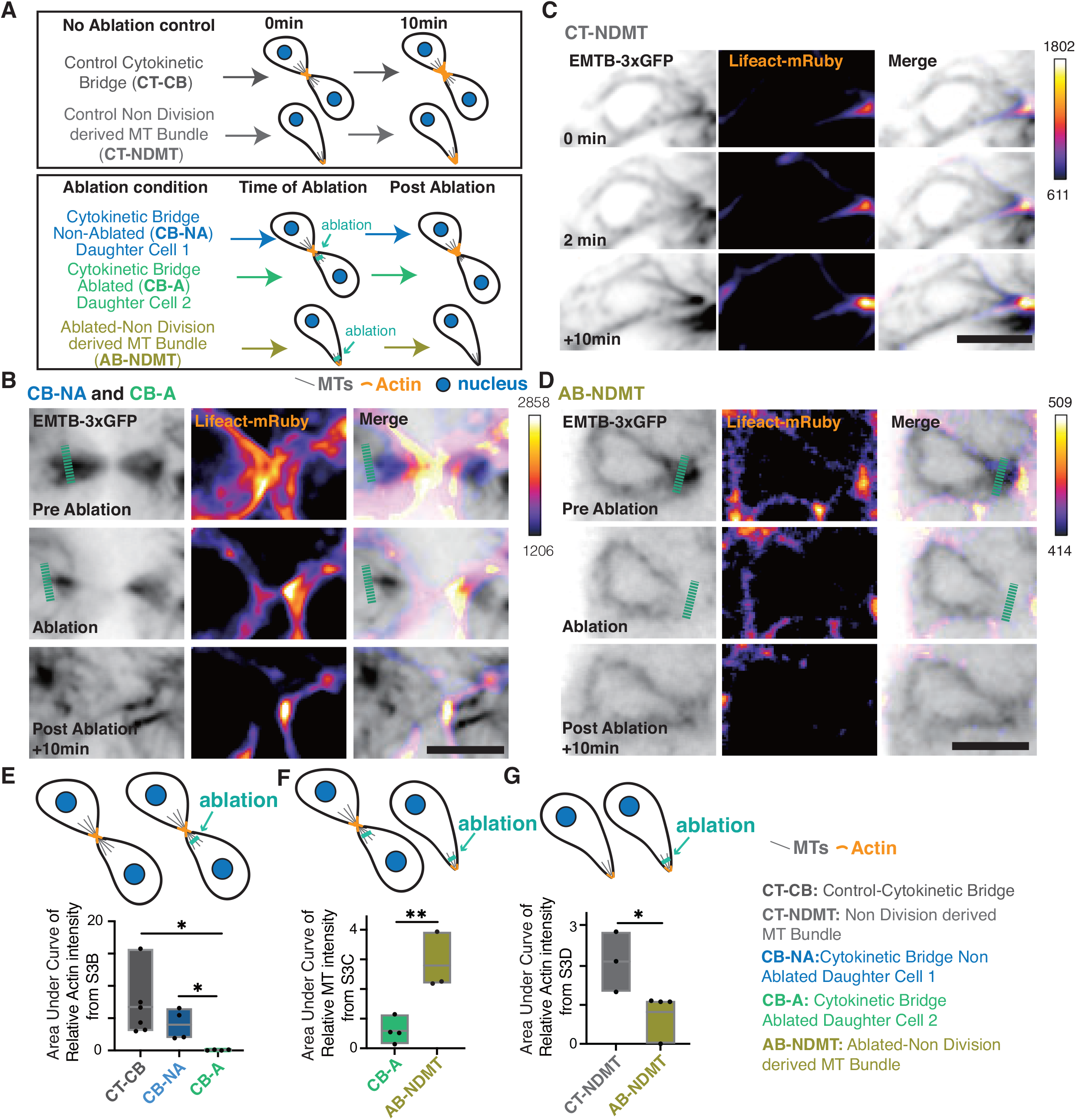
Cytokinetic bridge MT bundles are required for actin recruitment at rosette during lumen formation. **(A)** Model depicting laser ablation conditions during KV development. **(B-D)** Volumetric projections from video acquisition of an abscission event. Actin is labeled with Lifeact-mRuby (Fire-LUT), and MT are labeled with EMTB-3xGFP (Gray). Scale bar, 10 μm. Ablation conditions are noted above image and outlined in (**A**). **(E-G)** Area under the curve of relative actin (**E,G**) or MT intensity (**F**) over 10 min from **Fig S3A-C**. One-way ANOVA with Dunnett’s multiple comparisons, *p<0.05, n≥4 cell, n≥2 clutches per condition for (**E**). Unpaired t-test, *p<0.05, **p<0.01. n≥3 cell, n=2 clutches per condition for (**F, G**). Statistical results detailed in **Table S1**.

## Discussion

Studies of LRO formation have elucidated the fundamental role of LRO progenitors in forming the rosette-like structure with epithelial-like characteristics before the initiation of lumen formation ^8,15,38^. However, the steps preceding lumen formation is not clear, especially in the context of a living embryo. By studying KV formation in zebrafish embryos with novel transgenic and cell biological tools, our results suggest a model in which FGF-mediated signals induce the proliferation of a subset of early KV precursor cells, predominantly located anteriorly to the developing KV. Their progeny is then incorporated into the rosette before lumen formation, whereas the progeny from the later cell divisions were excluded (**Fig 8**). How FGF signaling coordinates the proliferation of the early KV precursors is an important question that awaits future studies.

**Figure 8:**
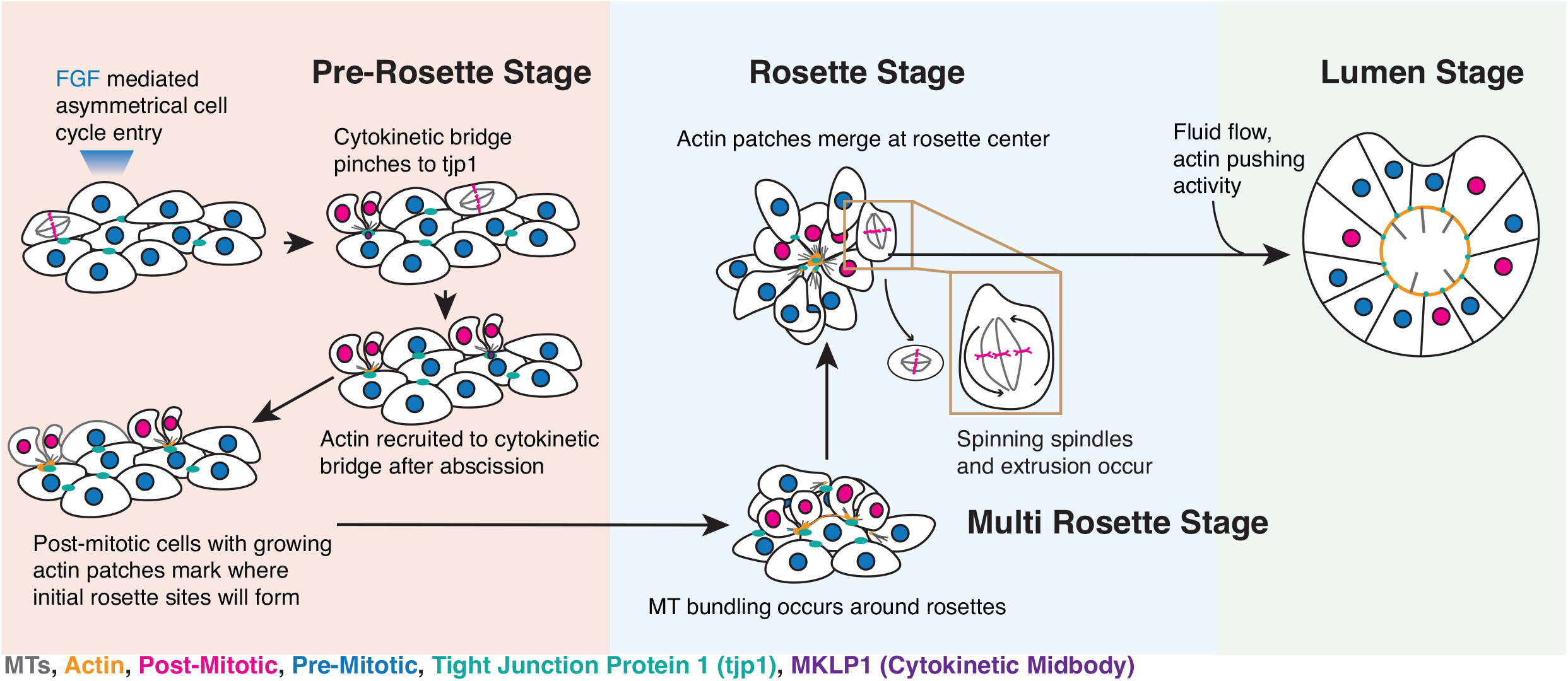
Proposed model for cell division contribution to lumen formation. Model depicting lumen formation through FGF mediated asymmetrical cell cycle entry which drives formation of cytokinetic bridge derived MT bundles that recruit apical actin patches at rosette centers. MT (cyan), actin (orange), post-mitotic nuclei (magenta), pre-mitotic nuclei (dark blue), tight junction protein 1 (tjp1, spring green), and midbody (MKLP1, purple) are shown.

We find that the apical microtubule (MT) bundles, which originate from the daughter cells of early Kupffer’s Vesicle (KV) precursors, persist after cytokinesis and are oriented towards regions marked by tjp1-labeled tight junctions (**Fig 8**). These apical MT bundles are crucial for recruiting actin to these sites. Actin recruitment has shown to be a process essential for apical membrane formation and lumen development, not only in KV but also in other systems such as mouse node formation ^8^ and pluripotent stem cell-derived epiblasts ^39^. For instance, in pluripotent stem cell-derived epiblasts, apical actin expansion is required for lumen formation, with subsequent lumen expansion driven by osmotic pressure-induced fluid transfer^39^. Under conditions of early mitotic ablation in KV where there is a significant reduction of actin to forming rosette structures, we observe that although a lumen forms, its size is significantly reduced (**Fig 3**). One possible explanation is that defects in tight junction assembly may lead to fluid leakage ^39–43^. However, our examination of lumen morphology in early mitotic-ablated KVs did not reveal the folded apical membrane appearance characteristic of such defects ^43^. This suggests that tight junction protein recruitment at the apical membrane in early mitotic-ablated KVs may be functional and formation occurs independently of mitotic events during lumen development. This is consistent with our observation that small tjp1-labeled tight junction structures are present before early mitotic events, with cytokinetic bridges pinching towards these structures (**Fig 8**). Nonetheless, the question remains whether osmolarity-driven fluid flow is sufficient to drive lumen opening under actin-deficient conditions and how lumen formation is initiated in such contexts.

In summary, our analysis of KV organization offers new insights into how spindle orientation, followed by mitotic exit driving MT bundle formation, contributes to actin recruitment during rosette formation, essential for KV development.

## Supporting information

Key Resource Table

## Acknowledgements

We thank SU’s BioImaging Core for their assistance and In Vivo Biosystems for helping us create the Sox17:EMTB-3xGFP line. This work was supported by National Institutes of Health Grants no. R01GM-127621 (H.H.), no. R01GM-130874 (H.H.), and no. OD026946 (SU BioImaging Center) for the Zeiss LSM980 with Airyscan2.

## Author Contribution

Y.W. and Y.L designed, performed, and analyzed experiments, and wrote the manuscript; H.H. oversaw the project and edited and wrote the manuscript; L.J. edited manuscript, and advised in animal studies; F.O., A.P., K.R., J.D.S., and A.W.S. performed experiments and associated analysis and edited the manuscript.

## Disclosure and competing interest statement

The authors declare no competing interests.

## Supplemental Figure Legends

**Figure S1.** Identification of an anteriorly positioned and pre-lumen enriched Kupffer’s Vesicle (KV) mitotic events and their post-mitotic distribution map.

*Related to **Fig 1***.

**(A)** KV mitotic cell position was mapped during KV development from **Video S1**. Different colors indicate different 20 min time bins pre and post lumen formation.

**(B)** Time bins from (**A**) and **Video S1** were mapped as percentage of mitotic KV cells in relation to lumen formation.

**(C, D)** Bubble plot depicting the positioning of mitotic event (PH3, **C**) and post mitotic cells (BrdU-positive, **D**) in KV at 8 and 12 hpf respectively. Large circle, represents > 7 events. Small circle represents 3 events. Tiny circle represents 1 event. n=26 embryos, across 3 clutches.

**(E)** 2 mitotic photoconversion events shown in **Fig 1F** highlighted. Scale bar, 10 μm. Statistical results detailed in **Table S1**.

**Figure S2.** An anterior/posterior FGF signaling gradient is required for anterior zones of mitotic activity. Related to **Fig 2**.

**(A-C)** Confocal projections of post lumen (12 hpf) embryos treated with methanol, 50 μM Cyclopamine (**A**), DMSO, 10 μM XAV939 (**B**), and 0.5μM, 1μM, 3μM, 5μM infigratinib (**C**). KV cell membranes marked with Sox17:GFP-CAAX (inverted gray), lumens highlighted in gold. Scale bar, 10 μm.

**(D)** KV morphologies characterized following infigratinib treatment with varying concentrations at 12 hpf, n>25 embryos per condition across 2 clutches.

**(E)** KV cell number at pre-lumen (8hpf) and post-lumen (12 hpf) following DMSO, 10 μM XAV939, and 1 μM infigratinib treatment. KV cell number was compared between *pcnt^+/+^* and *pcnt^-/-^* zebrafish. n>16 embryos (small dots represent embryos) across n≥3 clutches (large dots represent clutch) per condition. Graphs are scatter plots with mean of clutches ±SEM shown, n≥3 clutches. **p<0.01, ***p<0.005, ****p<0.0001, ns, not significant.

Statistical results detailed in **Table S1.**

**Figure S3.** Cytokinetic bridge MT bundles are required for actin recruitment at rosette during lumen formation. Related to Fig 7.

**(A)** Model depicting laser ablation conditions during KV development.

**(B-D)** Actin intensity (Lifeact-mRuby, **S3B, D**) and MT (EMTB-3xGFP, **S3C**) was measured over 10 min for cytokinetic bridge ablation conditions (**S3B, C**) and non-division-derived MT bundle (**S3C, D**) ablation conditions outlined in (**S3A**) and images in **Fig 7B-D**. n≥3±SEM.

## Supplemental Table Legends

**Table S1.** *Detailed statistical analysis of results reported in this study.* This table includes detailed result of all statistical analysis used. Figure where analysis applied, category of analysis, embryo sample number, number of clutches, statistic test applied, parameter, significant level, and specific p-values are shown.

## Supplemental Video Legends

**Video S1.** Identification of an anteriorly positioned and pre-lumen enriched Kupffer’s Vesicle (KV) mitotic events and their post-mitotic distribution map. Related to **Fig 1B**. Live confocal registered video showing mitotic events within KV in an embryo from the outcross of Sox17:GFP-CAAX and H2afx:h2afv-mCherry transgenic lines. KV cell plasma membranes marked by magenta (Sox17: GFP-CAAX) and nuclei (cyan, H2afx: h2afv-mCherry). Scale bar, 20 μm.

**Video S2.** Early KV mitotic events hold greater significance to KV development compared to later KV mitotic events. Related to **Fig 3C, D**.

Live confocal video showing lumen formation from a Sox17:GFP-CAAX, H2afx:h2afv-mCherry embryos under different laser ablation conditions. Sox17:GFP-CAAX (inverted gray) shown. Scale bar, 10 μm.

**Video S3:** KV spindles stably align along KVs longest axis until the KV starts rounding, then spindles spin and are extruded. Related to **Fig 4A, B**.

Live confocal video showing MT (grey, Sox17:EMTB-3xGFP) in KV (left). Highlighted from KV video on left is a spinning spindle (middle) and cell extrusion event (right). Scale bar, 20 μm.

**Video S4:** KV spindles stably align along KVs longest axis until the KV starts rounding, then spindles spin and are extruded. Related to **Fig 4G**.

Time lapse data set of Sox17:EMTB-3xGFP (MTs, Fire LUT) embryos under control (no ablation) conditions compared to laser ablation conditions (cond.1, first four mitotic events ablated; and cond. 3, all mitotic events ablated after KV reaches 20 cells). Scale bar, 20 μm.

**Video S5**: *Converging cytokinetic cells initiate actin recruitment at forming rosette centers*. *Related to* ***Fig 6A-6B***.

Time lapse of actin (Lifeact-mRuby, fire-LUT) and KV MTs (Sox17:EMTB-3xGFP, shown in inverted gray) during KV rosette formation. Scale bar, 10 μm.

## Resource Availability

### Lead contact

For further information or to request resources/reagents, contact Lead Contact, Dr. Heidi Hehnly

### Materials availability

New materials generated for this study are available for distribution.

### Data and code availability

All data sets analyzed for this study are displayed.

## Fish Lines

Zebrafish lines were maintained in accordance with protocols approved by the Institutional Animal Care Committee of Syracuse University (IACUC Protocol #18-006). Embryos were raised at 28.5°C and staged (as described in ^44^). Wildtype and/or transgenic zebrafish lines used for live imaging and immunohistochemistry are listed in Key Resource Table.

## Method Details

### Chemical inhibitors

Dechorionated live embryos were treated with 10 μM XAV939 (Sigma-Aldrich), 0.5 to 5 μM Infigratinib (MedChem Express), and 50 μM cyclopamine (Sigma-Aldrich) were prepared. Tg(sox17:GFP-CAAX) transgenic zebrafish embryos were dechorionated on 2% agarose pad dishes and at 50% epiboly incubated with vehicle control (0.4% DMSO, 0.5 % Methanol) and in listed concentrations of inhibitor diluted in zebrafish embryo water (0.03% sea salt, 1*10^-5^% Methylene Blue, RO water) and also taking into account the volume of agarose pad. Embryos were incubated to 8 hours post fertilization (hpf) and 12 hpf and fixed in 4% paraformaldehyde (PFA) (refer to immunofluorescence section).

### BrdU Incorporation

Tg(sox17:GFP-CAAX) embryos were dechorionated at 50% epiboly, followed by incubation in E3 buffer (5mM NaCl,0.17mM KCl, 0.33mM CaCl_2_,0.33mM MgSO_4_) for 15 minutes at room temperature. Embryos were then incubated in 10 mM BrdU in E3 buffer containing 12% DMSO and maintained at temperatures between 28.5 and 30°C until 10 hpf. Following serial dilution washes in E3 medium, the embryos were incubated until 12 hpf and fixed in 4% PFA and overnight dehydration in cold methanol at -20°C. Embryos then underwent a sequential rehydration process using methanol to PBST mixture ratios of 3:1, 1:1, and 1:3 for 5 minutes each step. Embryos were refixed for 15 minutes in 4% PFA with 0.5% Triton-X 100, followed by acid treatment with 2N HCl for 1 hour and subsequent neutralization in 0.1M borate buffer (0.1M Boric Acid, 0.076M NaOH) for 20 minutes. Then standard immunofluorescent approaches were used to label for BrdU (anti-BrdU, mouse, 1:100, Sigma-Aldrich,11170376001: BMC9318). See immunofluorescence section and Key Resource Table.

### Plasmids and mRNA for injection experiments

Plasmids were generated using Gibson cloning methods (NEBuilder HiFi DNA assembly Cloning Kit) and maxi-prepped before injection and/or transfection. mRNA was made using mMESSAGE mMACHINE^TM^SP6 transcription kit. See key resource table for a list of plasmid constructs and mRNA used. Injections of 1 cell staged embryos were performed as described in ^45^.

### Immunofluorescence

Immunostaining for phospho-histone H3 (PH3), acetylated tubulin and GFP, Tg(sox17:GFP-CAAX) transgenic zebrafish embryos were fixed at 8 and 12 hours post fertilization (hpf) using 4% PFA with 0.5% Triton X-100 overnight at 4°C. Embryos were dechorionated after washing with PBST (0.1% Tween-20 in Phosphate Buffered Saline (PBS)) 3 times. Embryos were blocked in wash solution (1%DMSO,1% BSA,0.1%Triton-X) for 1 hour at room temperature with gentile agitation. Primary antibody incubation (diluted in wash solution) occurs overnight at 4°C. Primary antibodies used include: anti-Phospho-Histone H3 (Rabbit) antibody (1:200, Cell Signaling Technology, 9701S), Anti-GFP (Chicken) (1:300, GeneTex, GTX13970: AB_371416) or Anti-Acetylated Tubulin (Mouse) (1:300, Sigma Aldrich,T6793: RRID: AB_477585), Anti-GFP (Rabbit) (1:300, Molecular Probes, A-11122: AB_221569), refer to key resource table. Embryos were then washed and incubated with secondary antibodies for 2-4 hours at room temperatures or overnight at 4°C. Secondary antibodies used include: Alexa Fluor Anti-Mouse 568 (1:300, Life Technologies, A10037; RRID: AB_2534013), Alexa Fluor Anti-Chicken 488 (1:300, Fisher scientific, A11039), or Alexa Fluor Anti-Mouse 647 (1:300, Life Technologies, A31571; RRID: AB_162542), Alexa Fluor Anti-Rabbit 488 (1:300, Life Technologies, A21206; RRID: AB_2535792). Embryos were stained with DAPI (1 μg/mL) to label nuclei after 3 times washes with wash solution. Embryos were mounted by 2% agarose after washing with PBS. Refer to Key resource table.

### Imaging

Fixed or live dechorionated embryos are embedded in low-melting 1.5% agarose (Key resource table) with the KV positioned at the bottom of a #1.5 glass bottom MatTek plate (Key resource table) and imaged using a spinning disk confocal microscope or laser scanning confocal microscope. Zebrafish embryos were imaged using Leica DMi8 (Leica, Bannockburn, IL) equipped with a X-light V2 Confocal unit spinning disk equipped with a Visitron VisiFRAP-DC photokinetics attached to 405 and 355nm lasers, a Leica SP8 (Leica, Bannockburn, IL) laser scanner confocal microscope (LSCM) and/or a Zeiss LSCM 980 (Carl Zeiss, Germany) with an Airyscan 2 detector. The Leica DMi8 is equipped with a Lumencore SPECTRA X (Lumencore, Beaverton, OR), Photometrics Prime-95B sCMOS Camera, and 89 North-LDi laser launch. VisiView software was used to acquire images. Optics used with this unit are HC PL APO x40/1.10W CORR CS2 0.65 water immersion objective, HC PL APO x40/0.95 NA CORR dry and HCX PL APO x63/1.40-0.06 NA oil objective. The SP8 laser scanning confocal microscope is equipped with HC PL APO 20x/0.75 IMM CORR CS2 objective, HC PL APO 40x/1.10 W CORR CS2 0.65 water objective and HC PL APO x63/1.3 Glyc CORR CS2 glycerol objective. LAS-X software was used to acquire images. The Zeiss LSM 980 is equipped with a T-PMT, GaASP detector, MA-PMT, Airyscan 2 multiplex with 4Y and 8Y. Optics used with this unit are PL APO x63/1.4 NA oil DIC. Zeiss Zen 3.2 was used to acquire the images. A Leica M165 FC stereomicroscope equipped with DFC 9000 GT sCMOS camera was used for staging and phenotypic analysis of zebrafish embryos.

### Photoconversion

Tg(sox17:GFP) embryos were injected with 75ng of Dendra-H2B mRNA (see methods in ^45^). The embryos were incubated at 30°C until 8 hpf, dechorionated and embedded in 1.5% agarose. To photoconvert H2B-Dendra, 405 nm laser beam was applied to a Region of Interest (ROI) placed over a mitotic plane. The 405 nm laser was used at 8 mW power with 2 FRAP cycles at 50 ms/pixel dwell time. Time-lapse videos were captured using 470 nm and 555 nm lasers to capture green and red emission from non-photoconverted and photoconverted respectively. Z-stacks were acquired at z-stack with a 2 μm step size every 2 minutes until 12 hpf.

### Laser Ablation

Tg(sox17:GFP-CAAX; h2afx:h2afv-mCherry) or Tg(sox17:EMTB-3XGFP) dechorionated live embryos were imaged on a spinning disk with VisiView kinetics unit starting at 6 hpf.

#### Mitotic ablation

Time-lapse videos capturing GFP-CAAX and h2afx:h2afv-mCherry used 470 nm and 555 nm lasers respectively across a z-stack with a 2 μm step size every 2 minutes until 12 hpf. KV mitotic events were ablated using a 355 nm pulsed laser operating at 65% power. The laser was applied within a ROI over mitotic cell for 2 cycles of 50 ms/pixel. Non-mitotic ablation control involved ablating a region containing 3-8 interphase cells of the KV. Ablated embryos were fixed at 1hpf and stained for cilia (acetylated tubulin).

#### MT bundles ablation

Time-lapse videos capturing EMTB-3XGFP and Lifeact-mRuby used 470 nm and 555 nm lasers, respectively, across a z-stack with a 2 μm step size every 2 minutes until 12 hpf. MT bundles (cytokinetic bridge-derived and non-division-derived bridge) were ablated using a 355 nm pulsed laser operating at 65% power. The laser was applied within an ROI over the proximal end of the bundles for 1-3 cycles of 50 ms/pixel until a break in the bundle was observed live.

### Image and data analysis

Images were processed using FIJI/ImageJ. Graphs and statistical analysis were produced using Prism 9 software. Additional analysis of KV cells were performed using Bitplane IMARIS software. Videos were created using FIJI/ImageJ or IMARIS. All time-lapse video projections are registered using FIJI. For percentage of ciliated KV cells, the number of cells with cilia was counted and represented as a percentage over the total number of cells in the cyst forming tissue.

#### Mitotic index and Anterior vs Posterior mitotic event calculation

For mitotic index, the number of mitotic cells (PH3 positive) in the KV was divided by the total number of KV cells (DAPI and Sox17:GFP-CAAX positive), resulting in a percentage of mitotic cells out of the entire population. To calculate fraction of mitotic events in the anterior versus posterior regions, anterior and posterior KV mitotic cell number was divided by the total number of mitotic cells in the KV.

#### KV Aspect Ratio

KV aspect ratio was calculated from a video projection of Sox17:EMTB-3xGFP embryos from 7 to 12 hpf. Images were acquired at maximum 2-to-3-minute intervals. For each captured frame, the aspect ratio was calculated using FIJI/ImageJ. Longest axis over shortest axis was determined giving the aspect ratio over time. Data was presented by averaging aspect ratios across 10 bins for the duration of KV development.

#### Angular Velocity of the Spindle

The metaphase derived spindle angle relative to the KV’s longest axis was measured over time. To do this, a line was positioned along the longest KV axis in relation to a second line that passes through the two spindle poles and the longest axis of a KV mitotic spindle. The ImageJ/FIJI angle tool calculated the angle at which the two lines intersect providing a spindle angle (∠Spindle). This spindle angle was calculated from metaphase to anaphase completion. The angular velocity was measured by taking the difference of identified spindle angle across consecutive time point 1 (∠Spindle_T1_) and time point 2 (∠Spindle_T2_) and dividing the difference by time interval (T_1_-T_2_).

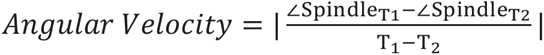

Data was presented by averaging spindle angular velocity from each mitotic event across 20 min bins for the duration of KV development between 7 to 12 hpf.

#### Lumen area

A region was drawn around the lumen perimeter and area calculated using the measure function. Where applicable, values were normalized to the control mean by dividing each lumen area by the mean value of the control lumens.

#### Actin intensity measurement at nascent MT bundles

KV cells of interest were masked across Z slices at five different time points using Imaris Software. A square ROI with constant size across all samples was placed at a newly formed MT bundle or MT bundle ablation site, labeled with EMTB-3xGFP, to quantify actin (Lifeact-mRuby) from max-projected datasets over time using the ROI manager in FIJI/ImageJ. Data normalization was performed by dividing the fluorescence intensity at each time point by the mean fluorescence intensity at the final time point for all measured bridges or intensity at the first time point after subtracting background intensity.

### Statistical Analysis

PRISM9 software was used for all graph preparations that include all individual data points across embryos and clutches denoted by color and size of points respectively and as noted in legends. These plots were presented as violin, floating bar, and a modified scatter plot that is termed “super-plot” ^46^ to denote individual embryos across clutches. Unpaired two-tailed t-tests for pairwise numerical differences in mean and one way ANOVA were performed when comparing greater than two groups using PRISM9 software. **** denotes a p-value<0.0001, *** p-value<0.001, **p-value<0.01, *p-value<0.05, ns, not significant. For further information on detailed statistical analysis see **Table S1**.

## Notes

### Competing Interest Statement

The authors have declared no competing interest.

### Summary of Updates

We have revised the abstract and added new findings from our original submission highlighted in Figure 5-7 and summarized in Figure 8.

## References Cited

1. Grimes, D.T., and Burdine, R.D. (2017). Left–Right Patterning: Breaking Symmetry to Asymmetric Morphogenesis. Trends in Genetics 33, 616–628. 10.1016/j.tig.2017.06.004.

2. Okabe, N., Xu, B., and Burdine, R.D. (2008). Fluid dynamics in zebrafish Kupffer’s vesicle. Dev Dyn 237, 3602–3612. 10.1002/DVDY.21730.

3. Dasgupta, A., and Amack, J.D. (2016). Cilia in vertebrate left-right patterning. Philos Trans R Soc Lond B Biol Sci 371. 10.1098/rstb.2015.0410.

4. Forrest, K., Barricella, A.C., Pohar, S.A., Hinman, A.M., and Amack, J.D. (2022). Understanding laterality disorders and the left-right organizer: Insights from zebrafish. Front Cell Dev Biol 10. 10.3389/FCELL.2022.1035513.

5. McGrath, J., Somlo, S., Makova, S., Tian, X., and Brueckner, M. (2003). Two populations of node monocilia initiate left-right asymmetry in the mouse. Cell 114, 61–73. 10.1016/S0092-8674(03)00511-7.

6. Layton, W.M. (1976). Random determination of a developmental process: reversal of normal visceral asymmetry in the mouse. J Hered 67, 336–338. 10.1093/OXFORDJOURNALS.JHERED.A108749.

7. Lee, J.D., and Anderson, K. V. (2008). Morphogenesis of the node and notochord: the cellular basis for the establishment and maintenance of left-right asymmetry in the mouse. Dev Dyn 237, 3464–3476. 10.1002/DVDY.21598.

8. Gredler, M.L., and Zallen, J.A. (2023). Multicellular rosettes link mesenchymal-epithelial transition to radial intercalation in the mouse axial mesoderm. Dev Cell 58, 933–950.e5. 10.1016/J.DEVCEL.2023.03.018.

9. Li, D., Mangan, A., Cicchini, L., Margolis, B., and Prekeris, R. (2014). FIP5 phosphorylation during mitosis regulates apical trafficking and lumenogenesis. EMBO Rep 15, 428–437. 10.1002/EMBR.201338128.

10. Mangan, A.J., Sietsema, D. V, Li, D., Moore, J.K., Citi, S., and Prekeris, R. (2016). Cingulin and actin mediate midbody-dependent apical lumen formation during polarization of epithelial cells. Nat Commun 7, 12426. 10.1038/ncomms12426.

11. Blasky, A.J., Mangan, A., and Prekeris, R. (2015). Polarized protein transport and lumen formation during epithelial tissue morphogenesis. Annu Rev Cell Dev Biol 31, 575–591. 10.1146/ANNUREV-CELLBIO-100814-125323.

12. Klinkert, K., Rocancourt, M., Houdusse, A., and Echard, A. (2016). Rab35 GTPase couples cell division with initiation of epithelial apico-basal polarity and lumen opening. Nat Commun 7. 10.1038/ncomms11166.

13. Boehlke, C., Kotsis, F., Buchholz, B., Powelske, C., Eckardt, K.U., Walz, G., Nitschke, R., and Kuehn, E.W. (2013). Kif3a guides microtubular dynamics, migration and lumen formation of MDCK cells. PLoS One 8. 10.1371/JOURNAL.PONE.0062165.

14. Mahjoub, M.R., and Stearns, T. (2012). Supernumerary Centrosomes Nucleate Extra Cilia and Compromise Primary Cilium Signaling. Current Biology 22, 1628–1634. 10.1016/J.CUB.2012.06.057.

15. Aljiboury, A.A., Ingram, E., Krishnan, N., Ononiwu, F., Pal, D., Manikas, J., Taveras, C., Hall, N.A., Da Silva, J., Freshour, J., et al. (2023). Rab8, Rab11, and Rab35 coordinate lumen and cilia formation during zebrafish left-right organizer development. PLoS Genet 19, e1010765. 10.1371/JOURNAL.PGEN.1010765.

16. Tavares, B., Jacinto, R., Sampaio, P., Pestana, S., Pinto, A., Vaz, A., Roxo-Rosa, M., Gardner, R., Lopes, T., Schilling, B., et al. (2017). Notch/Her12 signalling modulates, motile/immotile cilia ratio downstream of Foxj1a in zebrafish left-right organizer. Elife 6. 10.7554/ELIFE.25165.

17. Neugebauer, J.M., Amack, J.D., Peterson, A.G., Bisgrove, B.W., and Yost, H.J. (2009). FGF signalling during embryo development regulates cilia length in diverse epithelia. Nature 458, 651–654. 10.1038/NATURE07753.

18. Meyers, E.N., and Martin, G.R. (1999). Differences in left-right axis pathways in mouse and chick: functions of FGF8 and SHH. Science 285, 403–406. 10.1126/SCIENCE.285.5426.403.

19. Caron, A., Xu, X., and Lin, X. (2012). Wnt/β-catenin signaling directly regulates Foxj1 expression and ciliogenesis in zebrafish Kupffer’s vesicle. Development 139, 514–524. 10.1242/DEV.071746.

20. Sepulveda, G., Antkowiak, M., Brust-Mascher, I., Mahe, K., Ou, T., Castro, N.M., Christensen, L.N., Cheung, L., Yoon, D., Huang, B., et al. (2018). Co-translational protein targeting facilitates centrosomal recruitment of PCNT during centrosome maturation. Co-translational protein targeting facilitates centrosomal recruitment of PCNT during centrosome maturation in vertebrates 7, 241083. 10.1101/241083.

21. Mikule, K., Delaval, B., Kaldis, P., Jurcyzk, A., Hergert, P., and Doxsey, S. (2006). Loss of centrosome integrity induces p38—p53—p21-dependent G1— S arrest. Nature Cell Biology 2006 9:2 9, 160–170. 10.1038/ncb1529.

22. Purohit, A., Tynan, S.H., Valle, R., and Doxsey, S.J. (1999). Direct Interaction of Pericentrin with Cytoplasmic Dynein Light Intermediate Chain Contributes to Mitotic Spindle Organization. J Cell Biol 147, 481. 10.1083/JCB.147.3.481.

23. Chen, C.-T.T., Hehnly, H., Yu, Q., Farkas, D., Zheng, G., Redick, S.D., Hung, H.-F.F., Samtani, R., Jurczyk, A., Akbarian, S., et al. (2014). A Unique Set of Centrosome Proteins Requires Pericentrin for Spindle-Pole Localization and Spindle Orientation. Current Biology 24, 2327–2334. 10.1016/j.cub.2014.08.029.

24. Delaval, B., and Doxsey, S.J. (2010). Pericentrin in cellular function and disease. J Cell Biol 188, 181–190. 10.1083/JCB.200908114.

25. Zimmerman, W.C., Sillibourne, J., Rosa, J., and Doxsey, S.J. (2004). Mitosis-specific anchoring of γ tubulin complexes by pericentrin controls spindle organization and mitotic entry. Mol Biol Cell 15, 3642–3657. 10.1091/mbc.E03-11-0796.

26. Bergstralh, D.T., Dawney, N.S., and St Johnston, D. (2017). Spindle orientation: a question of complex positioning. Development 144, 1137–1145. 10.1242/DEV.140764.

27. Vertii, A., Kaufman, P.D., Hehnly, H., and Doxsey, S. (2018). New dimensions of asymmetric division in vertebrates. Preprint at John Wiley and Sons Inc., 10.1002/cm.21434 https://doi.org/10.1002/cm.21434.

28. Rathbun, L.I., Aljiboury, A.A., Bai, X., Hall, N.A., Manikas, J., Amack, J.D., Bembenek, J.N., and Hehnly, H. (2020). PLK1- and PLK4-Mediated Asymmetric Mitotic Centrosome Size and Positioning in the Early Zebrafish Embryo. Current Biology 30, 4519–4527.e3. 10.1016/j.cub.2020.08.074.

29. Gudipaty, S.A., and Rosenblatt, J. (2017). Epithelial Cell Extrusion: pathways and pathologies. Semin Cell Dev Biol 67, 132. 10.1016/J.SEMCDB.2016.05.010.

30. Marinari, E., Mehonic, A., Curran, S., Gale, J., Duke, T., and Baum, B. (2012). Live-cell delamination counterbalances epithelial growth to limit tissue overcrowding. Nature 484, 542–545. 10.1038/NATURE10984.

31. Nanavati, B.N., Yap, A.S., and Teo, J.L. (2020). Symmetry Breaking and Epithelial Cell Extrusion. Cells 9. 10.3390/CELLS9061416.

32. Eisenhoffer, G.T., and Rosenblatt, J. (2013). Bringing balance by force: live cell extrusion controls epithelial cell numbers. Trends Cell Biol 23, 185–192. 10.1016/J.TCB.2012.11.006.

33. Levic, D.S., Yamaguchi, N., Wang, S., Knaut, H., and Bagnat, M. (2021). Knock-in tagging in zebrafish facilitated by insertion into non-coding regions. Development 148. 10.1242/DEV.199994.

34. Steigemann, P., and Gerlich, D.W. (2009). Cytokinetic abscission: cellular dynamics at the midbody. Trends Cell Biol 19, 606–616. 10.1016/J.TCB.2009.07.008.

35. Steigemann, P., Wurzenberger, C., Schmitz, M.H.A., Held, M., Guizetti, J., Maar, S., and Gerlich, D.W. (2009). Aurora B-mediated abscission checkpoint protects against tetraploidization. Cell 136, 473–484. 10.1016/J.CELL.2008.12.020.

36. Guizetti, J., Schermelleh, L., Mäntler, J., Maar, S., Poser, I., Leonhardt, H., Müller-Reichert, T., and Gerlich, D.W. (2011). Cortical constriction during abscission involves helices of ESCRT-III-dependent filaments. Science (1979) 331, 1616–1620. 10.1126/SCIENCE.1201847/SUPPL_FILE/GUIZETTI.SOM.PDF.

37. Addi, C., Bai, J., and Echard, A. (2018). Actin, microtubule, septin and ESCRT filament remodeling during late steps of cytokinesis. Curr Opin Cell Biol 50, 27–34. 10.1016/J.CEB.2018.01.007.

38. Essner, J.J., Amack, J.D., Nyholm, M.K., Harris, E.B., and Yost, H.J. (2005). Kupffer’s vesicle is a ciliated organ of asymmetry in the zebrafish embryo that initiates left-right development of the brain, heart and gut. Development 132, 1247–1260. 10.1242/DEV.01663.

39. Indana, D., Zakharov, A., Lim, Y., Dunn, A.R., Bhutani, N., Shenoy, V.B., and Chaudhuri, O. (2024). Lumen expansion is initially driven by apical actin polymerization followed by osmotic pressure in a human epiblast model. Cell Stem Cell 31, 640–656.e8. 10.1016/J.STEM.2024.03.016.

40. Kuo, W.T., Odenwald, M.A., Turner, J.R., and Zuo, L. (2022). Tight junction proteins occludin and ZO-1 as regulators of epithelial proliferation and survival. Ann N Y Acad Sci 1514, 21–33. 10.1111/NYAS.14798.

41. Odenwald, M.A., Choi, W., Buckley, A., Shashikanth, N., Joseph, N.E., Wang, Y., Warren, M.H., Buschmann, M.M., Pavlyuk, R., Hildebrand, J., et al. (2017). ZO-1 interactions with F-actin and occludin direct epithelial polarization and single lumen specification in 3D culture. J Cell Sci 130, 243–259. 10.1242/JCS.188185.

42. Fanning, A.S., Van Itallie, C.M., and Anderson, J.M. (2012). Zonula occludens- 1 and -2 regulate apical cell structure and the zonula adherens cytoskeleton in polarized epithelia. Mol Biol Cell 23, 577–590. 10.1091/MBC.E11-09-0791/ASSET/IMAGES/LARGE/577FIG10.JPEG.

43. Mukenhirn, M., Wang, C.-H., Guyomar, T., Bovyn, M.J., Staddon, M.F., van der Veen, R.E., Maraspini, R., Lu, L., Martin-Lemaitre, C., Sano, M., et al. (2024). Tight junctions control lumen morphology via hydrostatic pressure and junctional tension. Dev Cell. 10.1016/J.DEVCEL.2024.07.016.

44. Kimmel, C.B., Ballard, W.W., Kimmel, S.R., Ullmann, B., and Schilling, T.F. (1995). Stages of embryonic development of the zebrafish. Developmental dynamics: an official public 203, 253–310. 10.1002/aja.1002030302.

45. Aljiboury, A.A., Mujcic, A., Cammerino, T., Rathbun, L.I., and Hehnly, H. (2021). Imaging the early zebrafish embryo centrosomes following injection of small-molecule inhibitors to understand spindle formation. STAR Protoc 2, 100293. 10.1016/j.xpro.2020.100293.

46. Lord, S.J., Velle, K.B., Dyche Mullins, R., and Fritz-Laylin, L.K. (2020). SuperPlots: Communicating reproducibility and variability in cell biology. Journal of Cell Biology 219. 10.1083/JCB.202001064/151717.

