## Supplemental Figures 1-3, Table S1 for "Specific Mitotic Events Drive Cytoskeletal Remodeling Required for Left-Right Organizer Development"

**Supplemental Information:**

Document S1. Figures S1-S3, and Table S1.

Video S1. *Identification of an anteriorly positioned and pre-lumen enriched Kupffer's Vesicle (KV) mitotic events and their post-mitotic distribution map.* Related to Fig 1B.

Video S2. *Early KV mitotic events hold greater significance to KV development compared to later KV mitotic events.* Related to Fig 3C, D.

Video S3. *Spindles stably align along KVs longest axis until the KV starts rounding, then spindles spin and are extruded.* Related to Fig 4A, B.

Video S4. *KV spindles stably align along KVs longest axis until the KV starts rounding, then spindles spin and are extruded.* Related to Fig 4G.

Video S5. *Converging cytokinetic cells initiate actin recruitment at forming rosette centers.* Related to Fig 6A-6B.

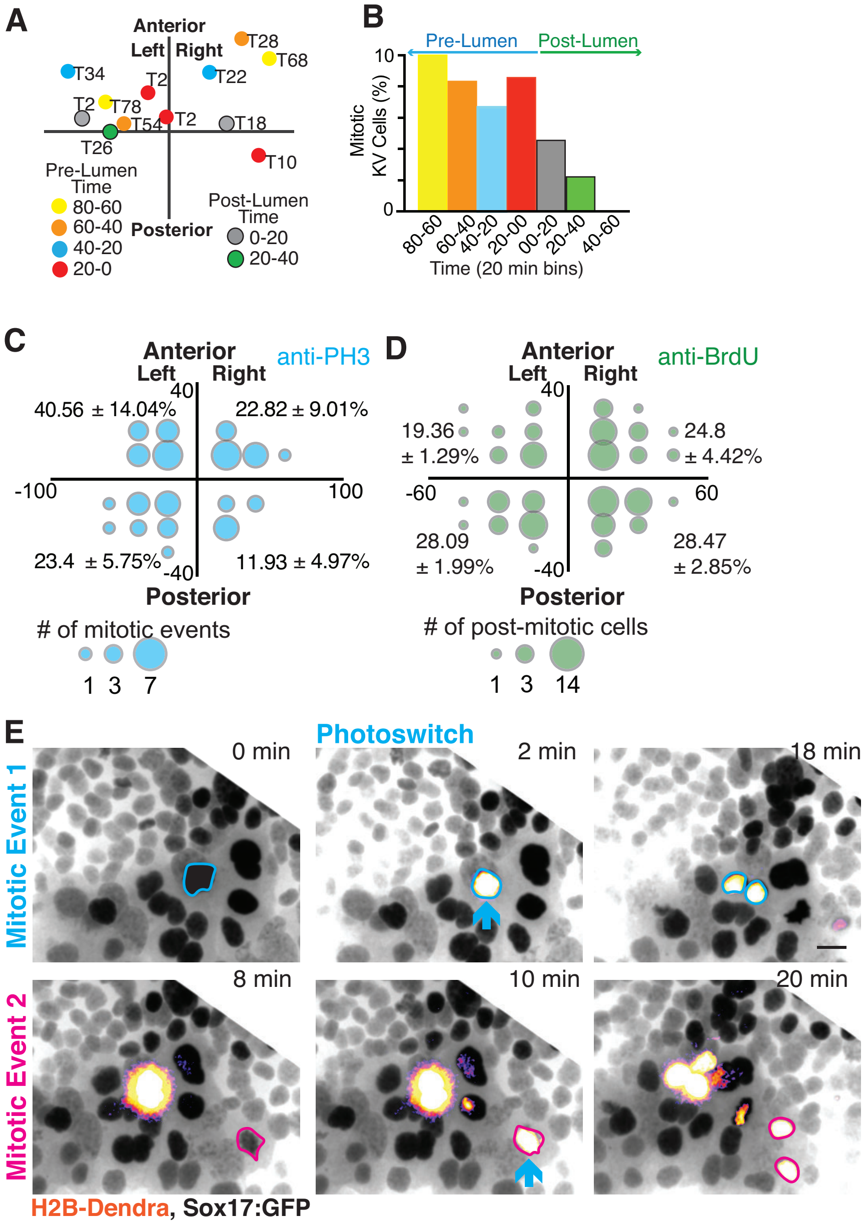


Fig S1. Identification of an anteriorly positioned and pre-lumen enriched KV mitotic events and their post-mitotic distribution map.

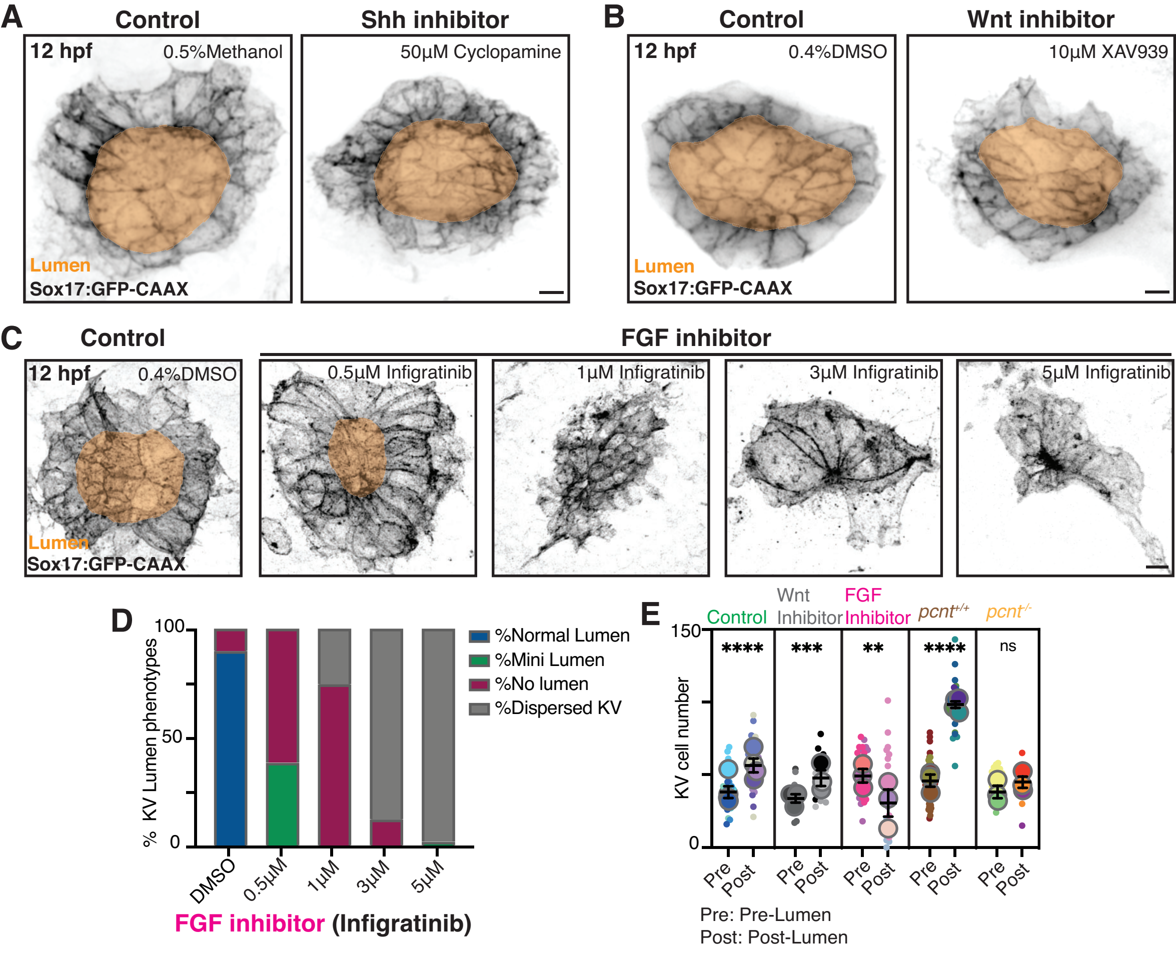


Fig S2. An anterior/posterior FGF signaling gradient is required for anterior zones of mitotic activity.

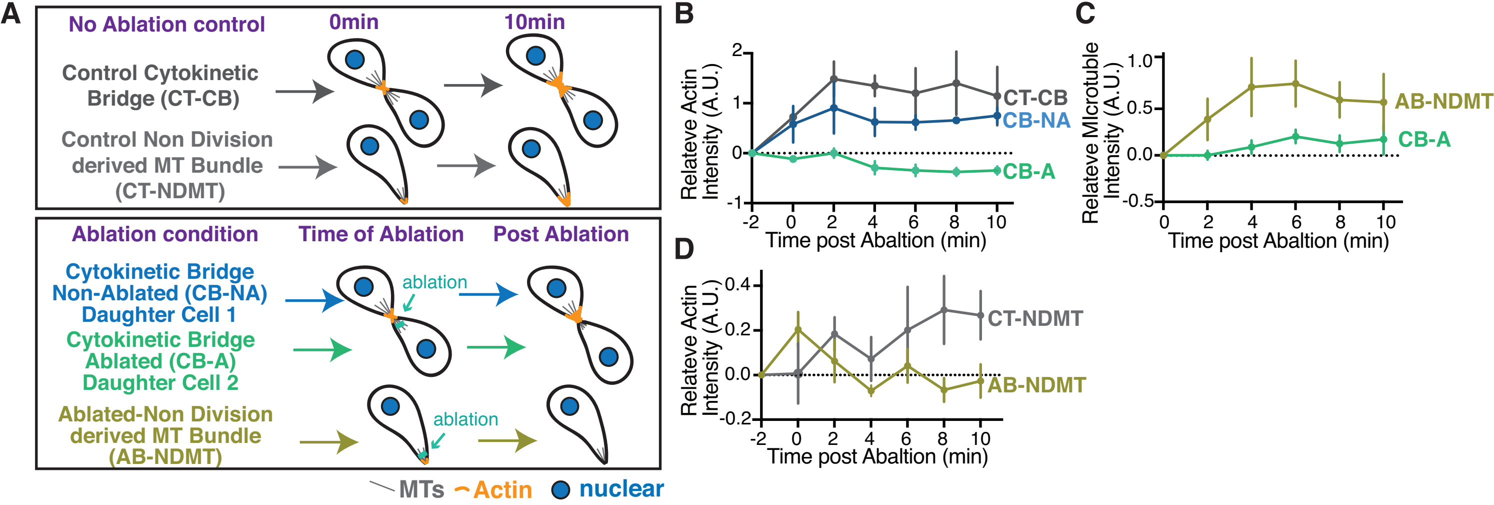


Fig S3. Cytokinetic bridge derived MT bundles are required for actin recruitment at rosette during lumen formation

**Table S1. Detailed statistical analysis of results reported in this study.**

| Figure | Category | n embryo | n Clutch | Statistical Test | Parameters | Result | p-value |
|---|---|---|---|---|---|---|---|
| **1C** | %Mitotic KV cells | 3 | 3 | N/A | N/A | N/A | N/A |
| **1E** | pH3 positive cell Anterior | 36 | 4 | Unpaired t-test | t=5.409, df=6 | N/A | N/A |
| | pH3 positive cell Posterior | | | | | ** | 0.0016 |
| | BrdU positive cell Anterior | 17 | 3 | Unpaired t-test | t=4.627, df=4 | N/A | N/A |
| | BrdU positive cell Posterior | | | | | ** | 0.0098 |
| **1I** | Ratio of Postmitotic anterior/posterior cell | 1 | 1 | N/A | N/A | N/A | N/A |
| **S1A** | %Mitotic KV cells | 1 | 1 | N/A | N/A | N/A | N/A |
| **S1B** | Position of the PH3 positive cell | 1 | 1 | N/A | N/A | N/A | N/A |
| **S1C** | pH3 positive cell | 26 | 3 | N/A | N/A | N/A | N/A |
| **S1D** | BrdU positive cell | 3 | 3 | N/A | N/A | N/A | N/A |
| **2D** | DMSO | 53 | 5 | One Way ANOVA (multiple comparison) | F (2,1.51) =5.621 | N/A | N/A |
| | XAV939 | 33 | 3 | | | ns | 0.4369 |
| | Infigratinib | 68 | 3 | | | ** | 0.0022 |
| | *pcnt* (+/+) | 46 | 3 | Unpaired t-test | t=2.638, df=89 | N/A | N/A |
| | *pcnt* (-/-) | 45 | 3 | | | ** | 0.0098 |
| **2E** | DMSO | 34 | 5 | Unpaired t-test | t=4.447, df=64 | **** | <0.0001 |
| | XAV939 | 20 | 3 | Unpaired t-test | t=2.262, df=38 | * | 0.0295 |
| | Infigratinib | 34 | 3 | Unpaired t-test | t=0.03508, df=66 | ns | 0.9721 |
| | *pcnt* (+/+) | 31 | 3 | Unpaired t-test | t=3.440, df=60 | ** | 0.0011 |
| | *pcnt* (-/-) | 30 | 3 | Unpaired t-test | t=2.199, df=58 | * | 0.0319 |
| **2F** | DMSO | 29 | 5 | One Way ANOVA | F (2,71) = 76.33 | N/A | N/A |
| | XAV939 | 16 | 3 | | | ns | 0.2 |

| | | | | | | | |
|---|---|---|---|---|---|---|---|
| | Infigratinib | 29 | 3 | (multiple comparison) | | **** | <0.0001 |
| | *pcnt* (+/+) | 16 | 4 | Unpaired t-test | t=6.018, df=57 | N/A | N/A |
| | *pcnt* (-/-) | 29 | 3 | | | **** | <0.0001 |
| **S2D** | DMSO | 39 | 2 | N/A | N/A | N/A | N/A |
| | 0.5 µM Infigratinib | 34 | 2 | N/A | N/A | N/A | N/A |
| | 1 µM Infigratinib | 39 | 2 | N/A | N/A | N/A | N/A |
| | 3 µM Infigratinib | 25 | 2 | N/A | N/A | N/A | N/A |
| | 5 µM Infigratinib | 57 | 2 | N/A | N/A | N/A | N/A |
| **S2E** | DMSO pre-lumen stage | 36 | 5 | Unpaired t-test | t=5.693, df=71 | N/A | N/A |
| | DMSO post-lumen stage | 37 | 4 | | | **** | <0.0001 |
| | XAV939 pre-lumen stage | 22 | 3 | Unpaired t-test | t=3.679, df=36 | N/A | N/A |
| | XAV939 post-lumen stage | 16 | 3 | | | *** | 0.0008 |
| | Infigratinib pre-lumen stage | 32 | 3 | Unpaired t-test | t=3.032, df=59 | N/A | N/A |
| | Infigratinib post-lumen stage | 29 | 3 | | | ** | 0.0036 |
| | *pcnt* (+/+) pre-lumen stage | 54 | 3 | Unpaired t-test | t=14.55, df=75 | N/A | N/A |
| | *pcnt* (+/+) post-lumen stage | 23 | 4 | | | **** | <0.0001 |
| | *pcnt* (-/-) pre-lumen stage | 35 | 3 | Unpaired t-test | t=1.906, df=56 | N/A | N/A |
| | *pcnt* (-/-) post-lumen stage | 23 | 3 | | | ns | 0.0617 |
| **3E** | No Ablation 8hpf | 4 | 4 | Unpaired t-test | t=3.586, df=7 | ** | 0.0089 |
| | No Ablation 12hpf | 5 | 5 | | | | |
| | Ablate All Mitotics 8hpf | 3 | 3 | Unpaired t-test | t=1.413, df=4 | ns | 0.2306 |
| | Ablate All Mitotics 12hpf | 3 | 3 | | | | |
| **3F** | Cond. 1_8hpf | 4 | 4 | Unpaired t-test | t=2.782, df=5 | * | 0.0388 |
| | Cond. 1_12hpf | 3 | 3 | | | | |

| | | | | | | | |
|---|---|---|---|---|---|---|---|
| | Cond. 2_8hpf | 3 | 3 | Unpaired t-test | t=4.230, df=4 | * | 0.0134 |
| | Cond. 2_12hpf | 3 | 3 | | | | |
| | Cond. 3_8hpf | 6 | 6 | Unpaired t-test | t=1.040, df=9 | ns | 0.3257 |
| | Cond. 3_12hpf | 5 | 5 | | | | |
| **3G** | Lumen area: No ablation control 20min | 3 | 3 | Unpaired t-test | t=1.000, df=4 | ns | 0.3739 |
| | Lumen area: All mitotic ablated 20min | 3 | 3 | | | | |
| | Lumen area: No ablation control 30min | 3 | 3 | Unpaired t-test | t=1.231, df=4 | ns | 0.2856 |
| | Lumen area: All mitotic ablated 30min | 3 | 3 | | | | |
| | Lumen area: No ablation control 40min | 3 | 3 | Unpaired t-test | t=1.356, df=4 | ns | 0.2466 |
| | Lumen area: All mitotic ablated 40min | 3 | 3 | | | | |
| | Lumen area: No ablation control 50min | 3 | 3 | Unpaired t-test | t=2.049, df=4 | ns | 0.1098 |
| | Lumen area: All mitotic ablated 50min | 3 | 3 | | | | |
| | Lumen area: No ablation control 60min | 3 | 3 | Unpaired t-test | t=1.777, df=4 | ns | 0.1503 |
| | Lumen area: All mitotic ablated 60min | 3 | 3 | | | | |
| | Lumen area: No ablation control 70min | 3 | 3 | Unpaired t-test | t=2.000, df=4 | ns | 0.1161 |
| | Lumen area: All mitotic ablated 70min | 3 | 3 | | | | |

| | | | | | | | |
|---|---|---|---|---|---|---|---|
| | Lumen area: No ablation control 80min | 3 | 3 | Unpaired t-test | t=2.174, df=4 | ns | 0.0954 |
| | Lumen area: All mitotic ablated 80min | 3 | 3 | | | | |
| | Lumen area: No ablation control 90min | 3 | 3 | Unpaired t-test | t=3.096, df=4 | * | 0.0364 |
| | Lumen area: All mitotic ablated 90min | 3 | 3 | | | | |
| | Lumen area: No ablation control 100min | 3 | 3 | Unpaired t-test | t=3.298, df=4 | * | 0.0300 |
| | Lumen area: All mitotic ablated 100min | 3 | 3 | | | | |
| | Lumen area: No ablation control 110min | 3 | 3 | Unpaired t-test | t=4.597, df=4 | * | 0.0101 |
| | Lumen area: All mitotic ablated 110min | 3 | 3 | | | | |
| **3H** | Lumen area: Cond. 1_10min | 3 | 3 | One Way ANOVA (multiple comparison) | F (2, 6) = 0.5000 | N/A | N/A |
| | Lumen area: Cond. 2_10min | 3 | 3 | | | ns | 0.6180 |
| | Lumen area: Cond. 3_10min | 4 | 4 | | | ns | 0.6180 |
| | Lumen area: Cond. 1_20min | 3 | 3 | One Way ANOVA (multiple comparison) | F (2, 6) = 0.5000 | N/A | N/A |
| | Lumen area: Cond. 2_20min | 3 | 3 | | | ns | 0.6180 |
| | Lumen area: Cond. 3_20min | 4 | 4 | | | ns | 0.6180 |
| | Lumen area: Cond. 1_30min | 3 | 3 | One Way ANOVA (multiple comparison) | F (2, 6) = 0.7719 | N/A | N/A |
| | Lumen area: Cond. 2_30min | 3 | 3 | | | ns | 0.4955 |
| | Lumen area: Cond. 3_30min | 4 | 4 | | | ns | 0.4955 |
| | Lumen area: Cond. 1_40min | 3 | 3 | One Way ANOVA | F (2, 6) = 2.493 | N/A | N/A |

| | | | | | | | |
|---|---|---|---|---|---|---|---|
| | Lumen area: Cond. 2_40min | 3 | 3 | (multiple comparison) | | ns | 0.1697 |
| | Lumen area: Cond. 3_40min | 4 | 4 | | | ns | 0.1697 |
| | Lumen area: Cond. 1_50min | 3 | 3 | One Way ANOVA (multiple comparison) | F (2, 6) = 1.520 | N/A | N/A |
| | Lumen area: Cond. 2_50min | 3 | 3 | | | ns | 0.2945 |
| | Lumen area: Cond. 3_50min | 4 | 4 | | | ns | 0.2945 |
| | Lumen area: Cond. 1_60min | 3 | 3 | One Way ANOVA (multiple comparison) | F (2, 6) = 8.375 | N/A | N/A |
| | Lumen area: Cond. 2_60min | 3 | 3 | | | * | 0.0215 |
| | Lumen area: Cond. 3_60min | 4 | 4 | | | * | 0.0215 |
| | Lumen area: Cond. 1_70min | 3 | 3 | One Way ANOVA (multiple comparison) | F (2, 6) = 19.10 | N/A | N/A |
| | Lumen area: Cond. 2_70min | 3 | 3 | | | ** | 0.0031 |
| | Lumen area: Cond. 3_70min | 4 | 4 | | | ** | 0.0031 |
| | Lumen area: Cond. 1_80min | 3 | 3 | One Way ANOVA (multiple comparison) | F (2, 6) = 48.23 | N/A | N/A |
| | Lumen area: Cond. 2_80min | 3 | 3 | | | *** | 0.0003 |
| | Lumen area: Cond. 3_80min | 4 | 4 | | | *** | 0.0003 |
| | Lumen area: Cond. 1_90min | 3 | 3 | One Way ANOVA (multiple comparison) | F (2, 6) = 17.48 | N/A | N/A |
| | Lumen area: Cond. 2_90min | 3 | 3 | | | ** | 0.0039 |
| | Lumen area: Cond. 3_90min | 4 | 4 | | | ** | 0.0039 |
| | Lumen area: Cond. 1_100min | 3 | 3 | One Way ANOVA (multiple comparison) | F (2, 6) = 35.51 | N/A | N/A |
| | Lumen area: Cond. 2_100min | 3 | 3 | | | *** | 0.0006 |
| | Lumen area: Cond. 3_100min | 4 | 4 | | | *** | 0.0006 |
| | Lumen area: Cond. 1_110min | 3 | 3 | One Way ANOVA (multiple comparison) | F (2, 6) = 25.79 | N/A | N/A |
| | Lumen area: Cond. 2_110min | 3 | 3 | | | ** | 0.0016 |

| | | | | | | | |
|---|---|---|---|---|---|---|---|
| | Lumen area: Cond. 3_110min | 4 | 4 | | | ** | 0.0016 |
| **3I** | Non-mitotic Control | 10 | 6 | One Way ANOVA (multiple comparison) | F (3, 17) = 3.302 | N/A | N/A |
| | Ablate Mitotic Cond. 1 | 3 | 3 | | | * | 0.0493 |
| | Ablate Mitotic Cond.2 | 4 | 4 | | | ns | 0.7471 |
| | Ablate Mitotic Cond. 3 | 5 | 5 | | | ns | 0.0893 |
| **3J** | Non-mitotic Control | 10 | 6 | One Way ANOVA (multiple comparison) | F (3, 18) = 9.935 | N/A | N/A |
| | Ablate Mitotic Cond. 1 | 3 | 3 | | | *** | 0.0003 |
| | Ablate Mitotic Cond. 2 | 4 | 4 | | | ns | 0.6507 |
| | Ablate Mitotic Cond. 3 | 5 | 5 | | | ns | 0.5365 |
| **4D** | Aspect Ratio of KV | 7 | 6 | N/A | N/A | N/A | N/A |
| **4E** | Spindle Angular Velocity | 43 Mitotic Cells | 6 | N/A | N/A | N/A | N/A |
| **4E'** | A.R.>2 | 12 cells (7 embryos ) | 6 | Unpaired t-test | t=3.114, df=30 | N/A | 0.0040 |
| | A.R.<2 | 20 cells (7 embryos ) | 6 | | | ** | 0.6507 |
| **4F, F'** | Pre-Lumen | 7 | 6 | Unpaired t-test | t=2.381, df=12 | N/A | 0.0347 |
| | Post Lumen | 7 | 6 | | | * | |
| **4G** | No Ablation Control | 7 | 6 | Unpaired t-test | t=2.488, df=11 | N/A | 0.0301 |
| | Ablate Mitotics Cond. 1 and 3 | 6 | 6 | | | * | |
| **6F** | CT-CB | 5 | 2 | N/A | N/A | N/A | N/A |

| | | | | | | | |
|---|---|---|---|---|---|---|---|
| | CT-NDMT | 5 | 2 | N/A | N/A | N/A | N/A |
| | Cond.1-NDMT | 3 | 1 | N/A | N/A | N/A | N/A |
| **6G** | CT-CB | 5 | 2 | One-way ANOVA (Multiple comparison) | F(2,10)=8.989 | * | 0.0185 |
| | CT-NDMT | 5 | 2 | | | N/A | |
| | Cond.1-NDMT | 3 | 1 | | | ns | 0.4377 |
| **7E** | CT-CB | 6 | 3 | One-way ANOVA (Multiple comparison) | F (2, 11) = 4.477 | * | 0.022 |
| | CB-NA | 4 | 3 | | | * | 0.0257 |
| | CB-A | 4 | 3 | | | N/A | |
| **7F** | CB-A | 4 | 2 | Unpaired t-test | t=4.103, df=5 | N/A | 0.0093 |
| | AB-NDMT | 3 | 2 | | | ** | |
| **7G** | CT-NDMT | 3 | 2 | Unpaired t-test | t=2.623, df=5 | N/A | 0.00469 |
| | AB-NDMT | 4 | 2 | | | * | |
| **S3B** | CT-CB | 6 | 3 | N/A | N/A | N/A | N/A |
| | CB-NA | 4 | 3 | N/A | N/A | N/A | N/A |
| | CB-A | 4 | 2 | N/A | N/A | N/A | N/A |
| **S3C** | CB-A | 3 | 2 | N/A | N/A | N/A | N/A |
| | AB-NDMT | 4 | 2 | N/A | N/A | N/A | N/A |
| **S3D** | CT-NDMT | 3 | 2 | N/A | N/A | N/A | N/A |
| | AB-NDMT | 4 | 2 | N/A | N/A | N/A | N/A |
