## Supplementary material for "Specific Mitotic Events Drive Cytoskeletal Remodeling Required for Left-Right Organizer Development": Key Resource Table

**Key resources table**

| REAGENT or RESOURCE | SOURCE | IDENTIFIER |
|---|---|---|
| Antibodies | | |
| Acetylated Tubulin | Sigma Aldrich | T6793: RRID: AB_477585 |
| Anti-GFP (Chicken) | GeneTex | GTX13970: AB_371416 |
| Anti-GFP (Rabbit) | Molecular Probes | A-11122: AB_221569 |
| Anti-PH3 (Rabbit) | Cell Signaling Technology | 9701S |
| Anti-BrdU (Mouse) | Sigma-Aldrich | 11170376001: BMC9318 |
| Alexa Fluor Anti-Rabbit 488 | Life Technologies | A21206; RRID: AB_2535792 |
| Alexa Fluor Anti-Rabbit 568 | Life Technologies | A10042; RRID: AB_2534017 |
| Alexa Fluor Anti-Chicken 488 | Fisher scientific | A11039 |
| Alexa Fluor Anti-Mouse 568 | Life Technologies | A10037; RRID: AB_2534013 |
| Alexa Fluor Anti-Mouse 647 | Life Technologies | A31571; RRID: AB_162542 |
| Alexa Fluor® 647 Phalloidin | Cell Signaling Technology | 8940S |
| Bacterial and virus strains | | |
| N/A | | |
| Biological samples | | |
| N/A | | |
| Chemicals, peptides, and recombinant proteins | | |
| DAPI | Sigma Aldrich | D9542-10mg |
| Agarose | Thermo Fisher | 16520100 |
| BSA | Fisher Scientific | BP1600-100 |
| Dimethylsulphoxide | Fisher Scientific | BP231-100 |
| Paraformaldehyde | Fisher Scientific | O4042-500 |
| Phosphate Buffered Saline | Fisher Scientific | 10010023 |
| Triton X-100 | Fisher Scientific | BP151500 |
| Tween 20 | Thermo Fisher | BP337500 |
| Sodium Chloride | Fisher Scientific | BP358 |
| 5-Bromo-2'-deoxyuridine, ≥99% (HPLC) | Sigma-Aldrich | Product Number : B5002<br>CAS-No. : 59-14-3 |
| XAV939 | Sigma-Aldrich | Product Number : X3004<br>CAS-No. : 284028-89-3 |
| Infigratinib | MedChem Express | Catalog No. : HY-13311<br>CAS No. : 872511-34-7 |

| | | |
|---|---|---|
| Cyclopamine | Sigma-Aldrich | Product Number : PHL82510<br>CAS-No. : 4449-51-8 |
| Critical commercial assays | | |
| BIO BASIC Maxi Prep Kit | BIO BASIC | 9K-0060023 |
| NEBuilder HiFi DNA assembly Cloning Kit | New England BioLabs | E5520S |
| mMESSAGE mMACHINETMSP6 | Invitrogen | AM1340 |
| Deposited data | | |
| N/A | | |
| Experimental models: Cell lines | | |
| N/A | | |
| Experimental models: Organisms/strains | | |
| Zebrafish | Zebrafish International Resource Center | AB-Wildtype |
| Zebrafish | Heidi Hehnly Lab | Sox17:GFP-CAAX; h2afx:h2afv-mCherry |
| Zebrafish | Jeffrey Amack Lab | Tg (sox17:GFP-CAAX)sny101 |
| Zebrafish | Li-En Jao Lab | PCNT (-/-) |
| Zebrafish | Zebrafish International Resource Center | Tg(sox17:GFP) |
| Zebrafish | Heidi Hehnly lab | Tg(sox17:EMTB-3×GFP) |
| Zebrafish | Heidi Hehnly lab | sox17:GFP-CAAX;PCNT(-/-) |
| Zebrafish | Heidi Hehnly lab<br>Michel Bagnat lab | Sox17:GFP-CAAX; Tjp1atdTomato |
| Oligonucleotides | | |
| H2B-Dendra (mRNA) | Heidi Hehnly lab | Plasmid: pCS2-H2B-Dendra |
| Lifeact-mRuby (mRNA) | Heidi Hehnly lab | Plasmid: pCS2-Lifeact-mRuby |
| mKate-MKLP1 (mRNA) | Heidi Hehnly lab | Plasmid: pCS2-mKate-MKLP1 |
| Recombinant DNA | | |
| Plasmid: pCS2-H2B-Dendra | Addgene | Plasmid #140559 |
| Plasmid: pCS2-Lifeact-mRuby | Addgene | Plasmid #194339 |
| Plasmid: pCS2-mKate-MKLP1 | Addgene | Plasmid #140570 |
| Software and algorithms | | |
| ImageJ/FIJI | NIH and Laboratory for Optical and Computational Instrumentation | https://imagej.net/Fiji |
| IMARIS, Bitplane | Oxford Instruments | https://imaris.oxinst.com/ |
| PRISM9 | GraphPad | https://www.graphpad.com/scientific-software/prism/ |

| | | |
|---|---|---|
| LAS-X Software | Leica Microsystems | https://www.leica-microsystems.com/products/microscope-software/p/leica-las-x-ls/ |
| VisiView | Visitron | https://www.visitron.de/products/visiviewr-software.html |
| Other | | |
| N/A | | |
